# Spatial and Temporal Regulation of Parent-of-Origin Allelic Expression in the Endosperm

**DOI:** 10.1101/2022.01.29.478178

**Authors:** Yuri S. van Ekelenburg, Karina S. Hornslien, Tom Van Hautegem, Matyáš Fendrych, Gert Van Isterdael, Katrine N. Bjerkan, Jason R. Miller, Moritz K. Nowack, Paul E. Grini

**Author notes:** Author Contributions: PEG, MKN designed the research; YSvE, TvH, MF, KNB, GvI, MNK & PEG performed the experiments; YSvE, KSH, TvH, MF, JRM, MNK & PEG analyzed and discussed the data; YSvE, KSH & PEG wrote the article; All authors revised and approved the article.

## Abstract

Genomic imprinting promotes differential expression of parental alleles in the endosperm of flowering plants, and is regulated by epigenetic modification such as DNA methylation and histone tail modifications in chromatin. After fertilization, the endosperm develops through a syncytial stage before it cellularizes and becomes a nutrient source for the growing embryo. Both in early and late endosperm development regional compartmentalization has been shown, and different transcriptional domains suggest divergent spatial and temporal regional functions. The analysis of the role of parent-of-origin allelic expression in the endosperm as a whole and also investigation of domain specific functions has been hampered by the availability of the tissue for high-throughput transcriptome analyses and contamination from surrounding tissue. Here we have used Fluorescence-Activated Nuclear Sorting (FANS) of nuclear targeted eGFP fluorescent genetic markers to capture parental specific allelic expression from different developmental stages and specific endosperm domains. This RNASeq approach allows us to successfully identify differential genomic imprinting with temporal and spatial resolution. In a systematic approach we report temporal regulation of imprinted genes in the endosperm as well as region specific imprinting in endosperm domains. Our data identifies loci that are spatially differentially imprinted in one domain of the endosperm while biparentally expressed in other domains. This suggests that regulation of genomic imprinting is dynamic and challenges the canonical mechanisms for genomic imprinting.

## Introduction

The double fertilization event in angiosperms forms the diploid zygote and triploid endosperm, respectively (Nowack et al., 2010). Endosperm development is characterized by two different stages, a syncytial stage and a cellular stage, which have different functions (Berger et al., 2006). The cellular endosperm can be further divided into subregions, and it has become evident that such endosperm domains have distinct expression profiles (Brown et al., 1999; Belmonte et al., 2013; Del Toro-De León and Köhler, 2019; Picard et al., 2021).

Due to the homodiploid nature of the embryo-sac central cell, fertilization by a haploid sperm cell gives rise to an unbalanced endosperm in regards to parental allelic contribution (2:1 maternal:paternal), and therefore tight regulation is required for balanced parental gene expression (Birchler, 1993). The endosperm, analogous to the mammalian placenta, is the prime site for the epigenetic phenomenon genomic imprinting, parent-of-origin dependent expression of genes due to epigenetic marks (Feil and Berger, 2007). Traditionally, this process involves partial or full silencing of one of the parental alleles and is associated with both DNA methylation and histone modification (Jullien et al., 2006; Satyaki and Gehring, 2017). Furthermore, the involvement of small RNAs through the RNA-directed DNA methylation (RdDM) pathway resulting in *de novo* DNA methylation has been demonstrated (Vu et al., 2013; Hornslien et al., 2019; Satyaki and Gehring, 2019; Batista and Köhler, 2020).

The functional role of genomic imprinting in flowering plants has not been fully understood as most mutants of imprinted genes do not show an obvious seed phenotype (Shirzadi et al., 2011; Wolff et al., 2015; Zhang et al., 2018; Bjerkan et al., 2020). Moreover, the number of maternally and paternally imprinted genes previously identified remains controversial due to the detection of widespread contamination from the maternal seed coat and diploid embryo tissue (Schon and Nodine, 2017). Therefore, it is crucial for the study of genomic imprinting to avoid contamination from genetically distinct surrounding tissues. Furthermore, it has been shown that parent-of-origin specific expression can be specific to endosperm regions, as suggested by differences in parental expression bias between domains (Picard et al., 2021). Although some examples have been identified where imprinted gene expression is completely inactivated (Kirkbride et al., 2019) or becomes increasingly biparental (Ngo et al., 2012), dynamic regulation of imprinting patterns in a spatial or temporal manner has not been systematically investigated for the *Arabidopsis thaliana* endosperm.

Here, we aim to enhance our understanding of the role of genomic imprinting in flowering plants by investigating spatio-temporal dynamics of parent-of-origin allelic expression. Since different gene regulatory programs operate in different endosperm domains, specific mechanisms to modulate temporal and spatial specific imprinting may have evolved. We hypothesize that genomic imprinting in the *A. thaliana* endosperm is dynamically regulated in both a temporal and spatial manner. In order to avoid contamination issues that complicated previous imprinting studies, we have analyzed transcriptomes from isolated nuclei of endosperm specific domains. To this end, we have generated nuclear targeted eGFP fluorescent genetic markers that report different temporal developmental stages or have a spatial resolution in endosperm domains. Using Fluorescence-Activated Nuclear Sorting (FANS) captured nuclei, we have successfully identified imprinted genes with an imprinting profile specific to developmental stages as well as subdomains of the endosperm. Our findings suggest that regulation of genomic imprinting is dynamic, both in a spatial and temporal perspective, challenging the canonical mechanisms for genomic imprinting.

## Results and Discussion

### Generation of endosperm specific temporal and spatial expression marker lines

In order to investigate the temporal and spatial effects on parent-of-origin specific allelic expression in *A. thaliana,* we developed endosperm specific genetic reporters that had specific temporal and spatial profiles. A dual component system was used where nuclear-localized histone 2A-GFP fusion protein (H2A-GFP) is expressed under control of a domain specific promoter *(proMARKER>>H2A-GFP)* (Weijers et al., 2003; Olvera-Carrillo et al., 2015). Based on seed tissue microarray data (Le et al., 2010) and on gene expression patterns (Winter et al., 2007) 39 candidate endosperm specific gene promoters were selected and cloned expression vectors were transformed into *A. thaliana* Col-0 (STable 1). The spatial and temporal expression of these constructs was characterized during seed development in 48 independent T1 individuals per reporter line. For the purpose of fluorescence-activated nuclear sorting (FANS) four lines with domain or stage specific markers were selected: AT5G09370 expressing in Early Endosperm (EE); AT3G45130 expressing in Embryo-Surrounding Region (ESR); AT4G31060 expressing in Developing Aleurone Layer (DAL); and AT4G00220 expressing in Total Endosperm (TE1). For each line, the expression pattern was verified in subsequent generations by confocal imaging (Figure 1). No seed coat or embryo GFP signal was observed in any of these lines. To demonstrate the utilization of the marker lines to identify cellularized endosperm nuclei, we performed FANS using an optimized sorting protocol to separate GFP positive nuclei from GFP negative nuclei (SFigure 1A and B). The ploidy levels of GFP positive nuclei corresponded to endosperm ploidies (3C,6C) compared to GFP negative (2C, 4C and 8C) ploidies (SFigure 1C). Additionally, in order to verify the specificity of endosperm spatial markers, the level of GFP transcript from GFP positive and GFP negative ESR and DAL nuclei was assessed by real-time qPCR (RT-qPCR) (SFigure 1D), demonstrating a 50 – 250 fold GFP transcript enrichment in GFP positive nuclei compared to GFP negative nuclei.

**Figure 1.**
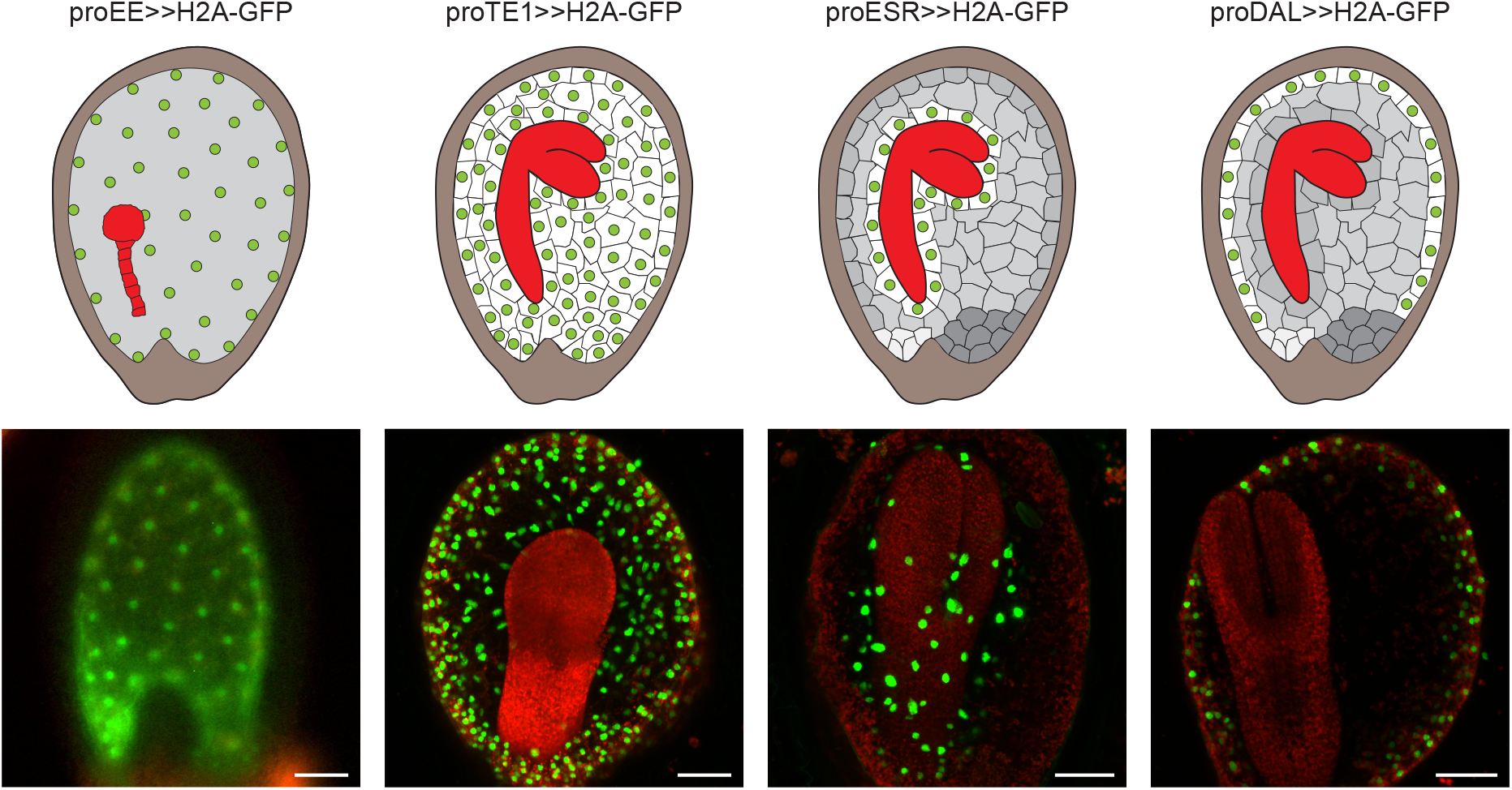
Verification of endosperm domain specific marker lines. Top panel, Cartoon representation of endosperm domains for the selected marker lines within the temporal and spatial plane. The GFP expressing endosperm nuclei for the selected marker lines are highlighted in green. The embryo and seed coat are represented in red and brown respectively. Endosperm that is not expressing GFP is depicted in grey. Bottom panel, Verification of domain specific GFP expression in endosperm domain marker lines using fluorescence microscopy (bottom). Red indicates autofluorescence of chloroplasts. proEE>>H2A-GFP = early endosperm; proTE1>>H2A-GFP = total endosperm; proESR>>H2A-GFP = embryo surrounding region; proDAL>>H2A-GFP = developing aleurone layer. Scale bar = 50 μm.

### Fluorescence-activated nuclear sorting of endosperm nuclei

In order to isolate endosperm domain-specific transcript profiles for temporal and spatial genomic imprinting analysis, homozygous marker lines EE, ESR, DAL and TE1 in the Col-0 accession were crossed as mothers to wild-type (WT) of the Tsu-1 accession. EE seeds were collected from dissected siliques at 4 days after pollination (DAP) and seeds from ESR, DAL and TE1 were collected at 7 DAP (SData 1). Seeds were homogenized to release nuclei and GFP positive nuclei were sorted by FANS. Nuclear RNA was isolated and cDNA libraries were sequenced yielding 150 bp paired-end reads (SData 1). Although nuclear RNA contains higher levels of pre-RNA and unspliced transcripts (Long et al., 2021), a high correlation between nuclear and total cellular mRNA has previously been demonstrated in plants, also including triploid endosperm tissue (Jacob et al., 2007; Deal and Henikoff, 2010; Del Toro-De León and Köhler, 2019; Picard et al., 2021). An average of 25 million reads per replicate mapped to Col-0 and Tsu-1 reference transcriptomes polished by Pilon (Walker et al., 2014) (SData 1).

To evaluate the variation between individual replicates of each marker line, reads were mapped separately to both the Col-0 and Tsu-1 polished transcriptomes. Counts from the two mappings were subjected to differential expression analysis (SData 2). Principal component analysis (PCA) of normalized read counts per gene explained 95% of the variances in the first two principal components. The PCA plot (Figure 2A) shows high between-group variation for all four lines, and low within-group variation for three lines. The analysis indicated high homogeneity of EE, TE1 and ESR replicates (Figure 2A), suggesting a high specificity of the sorting process. However, divergence in the DAL line (Figure 2A), combined with its insufficient replicates and read counts (SData 1), indicated the DAL domain marker data should be omitted from further analyses. To further address the sensitivity of the marker generated transcriptomes, the differential expression analysis was repeated excluding DAL and used to assess temporal (EE vs TE1), spatial (ESR vs TE1) and spatial-temporal (EE vs ESR) expression differences (SData 2). As expected, due to comparison of different tissues or developmental time-points, the differential expression was most pronounced when comparing both spatial and temporal expression (EE vs ESR – 8714 differentially expressed genes), followed by differential expression in 4 DAP vs 7 DAP stages (EE vs TE1 – 6979 differentially expressed genes) (SFigure 2A and B respectively). Importantly, albeit the embryo-surrounding region (ESR) is contained within the total endosperm (TE1), 5324 genes were also differentially regulated comparing ESR with TE1 at 7 DAP (Figure 2B). This demonstrates that significant expression changes between total endosperm and the embryo surrounding region can be visualized using profiles from TE1 and ESR FANS. We next explored whether allelic parent-of-origin specific expression could be addressed with these data.

**Figure 2.**
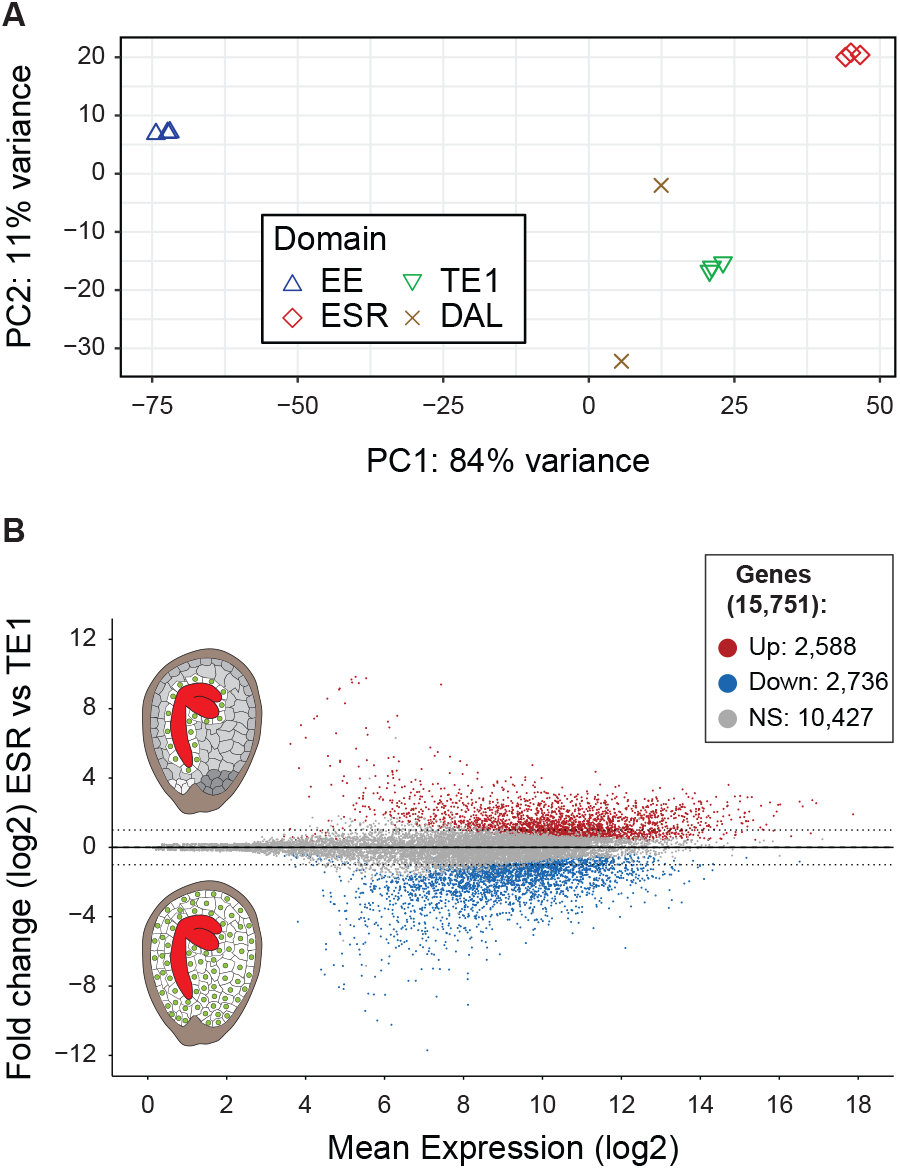
Endosperm marker lines homogeneity assessment and verification of spatial differential gene expression. A, Principal component analysis (PCA) of gene expression of individual biological replicates for the respective marker lines; EE (blue triangle), ESR (red diamond), DAL (brown cross) and TE1 (green triangle) to assess the homogeneity within replicates and between each marker line. B, MA plot showing the fold change (log2) differential expression between ESR and TE1 versus the mean expression (log2). The analysis revealed 5,324 genes to be significantly differentially expressed (red = up, blue = down, adjusted p-value < 0.05) within the spatial plane. Non significant (NS) genes are depicted in grey. Dashed lines indicate log2 of 1 and −1. EE = early endosperm; ESR = embryo surrounding region; DAL = developing aleurone layerTE1 = total endosperm.

### Informative read calling identifies imprinted genes

Prior to imprinting analysis various filters were applied (STable 2, SData 3) as described previously (Hornslien et al., 2019). Genes with a general expression bias between the ecotype accessions were identified by differential expression analysis of homozygous Col-0 and Tsu-1 samples and filtered (SFigure 4, SData 4). A read pair was considered informative if it mapped to the same gene in the polished transcripts from both parents, such that alignment features (InDels, SNPs) distinguished the parental allele of origin (Hornslien et al., 2019). Ultimately, 6169, 7847 and 8024 genes with informative reads were selected for allele-specific differential expression analysis in EE, ESR and TE1 respectively (STable 2, SData 5).

Genes were classified based on the total informative read count, parental bias, and statistical significance (SFigure 5). First, we identified genes with an insufficient number of informative read counts. Second, genes that did not show a parental bias were identified. Third, genes that showed non-significant parental bias were identified. Finally, genes showing a parental bias and a significant fold change (FC) were considered imprinted (SFigure 5).

In total, across all investigated marker lines, 181 maternally expressed genes (MEGs) and 56 paternally expressed genes (PEGs) were identified. The ratios of MEG:PEG per line was 82:28 in EE, 60:11 in TE1, 69:22 in ESR, and 181:56 overall (SData 6).Gene ontology (GO) enrichment analysis identified significantly enriched GO terms in the imprinted gene panels (STable 3, SData 6). Overall, enrichment was found for genes encoding proteins involved in gene regulation. For all identified gene panels, out of 237 genes that could be tested, transcription factor activity was enriched (1,8 fold, n=23). More specifically, all MEGs taken together (2 fold, n=20) and also all single MEG panels were enriched for transcription factor activity (STable 3, SData 6), including proteins from several transcription factor (TF) families, including Homeobox TFs *(FWA, BEL1, KNAT3, RPL),* MYB domain TFs *(MYB7, MYB60),* MADS-box *(STK),* WRKY *(WRK50),* heat shock *(HSF4),* zinc-finger *(ZF3),* EAR-containing *(TIE4)* and basic helix-loop-helix *(TT8)* proteins.

### Comparison of imprinting studies

In order to compare our data to previously reported imprinted gene sets, we estimated the overlap between previous recent studies and this study ((Pignatta et al., 2014; Del Toro-De León and Köhler, 2019; Hornslien et al., 2019; Picard et al., 2021); SData 7). There is measurable overlap of identified imprinted gene identities between our study and other recent studies (Figure 3). All FANS marker lines (EE, TE1, ESR) identified imprinted genes that had been reported previously. The largest overlap to previous studies was found with EE (38 MEGs and 13 PEGs). The overlap to previously identified imprinted genes identified using the ESR marker was low for paternally expressed genes (20 MEGs and two PEGs) suggesting that the ESR experiment may report imprinted genes masked by total endosperm in previous studies. All markers taken together, a total of 78 imprinted genes (60 MEGs and 18 PEGs; SData 8) out of 237 total imprinted genes were previously identified (33%), including *FLOWERING WAGENINGEN (FWA), SEEDSTICK (STK), BANYULS (BAN), FIDDLEHEAD (FDH), YUCCA10 (YUC10), HOMEODOMAIN GLABROUS 3 (HDG3) PEG3* and *PEG6* (Pignatta et al., 2014; Del Toro-De León and Köhler, 2019; Hornslien et al., 2019; Picard et al., 2021).

**Figure 3.**
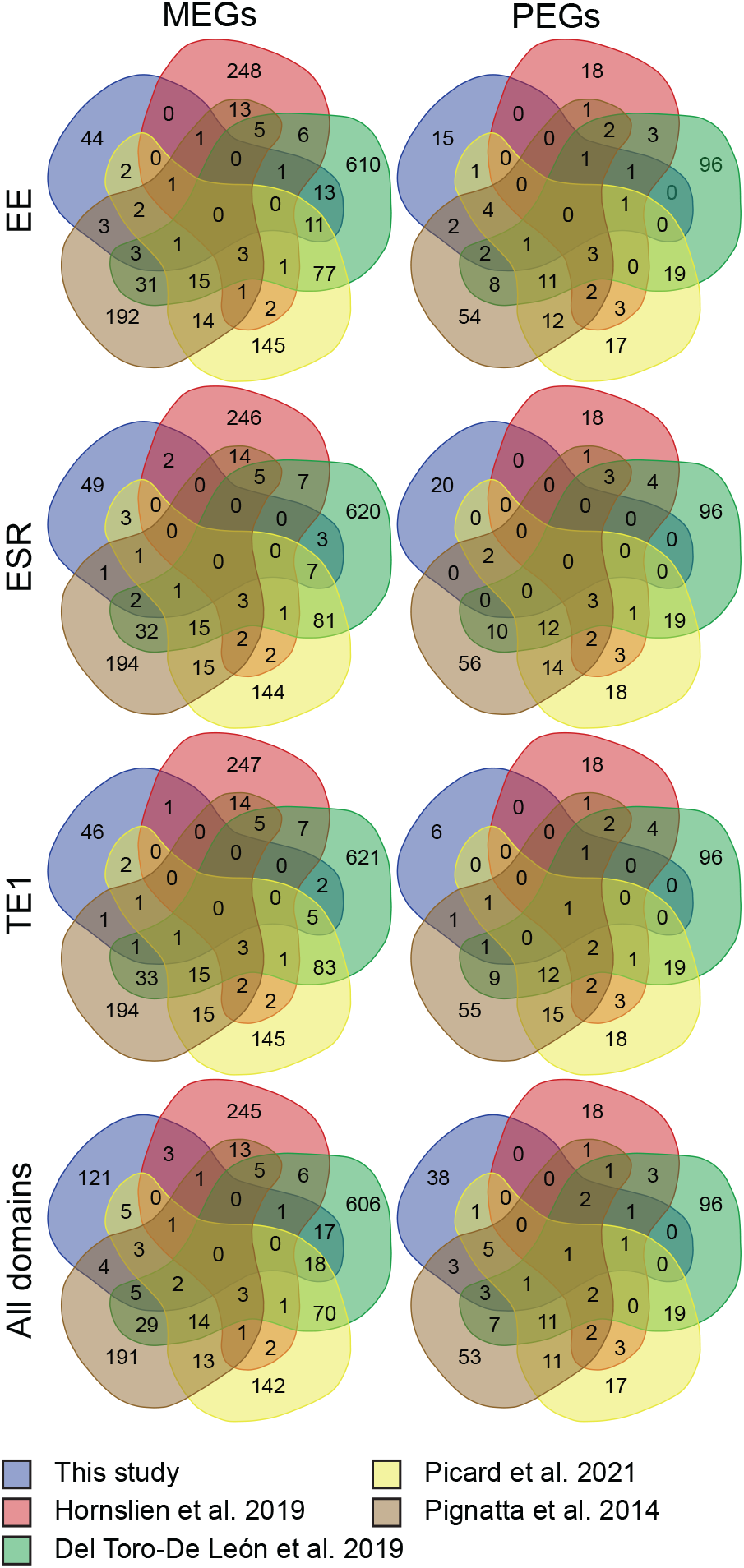
Identified MEGs and PEGs overlap with previous studies. Overlap of identified MEGs (left column) and PEGs (right column) for each domain separately (EE, ESR and TE1) and combined (All domains) between this study (blue) and previous studies (Pignatta et al., 2014) [brown], (Hornslien et al., 2019) [red], (Del Toro-De León and Köhler, 2019) [green] and (Picard et al., 2021) [yellow]).

The high degree of uniquely identified imprinted genes when contrasting our study with other reports as well as between these reports are likely explained by methodological difference and experimental setup. Most studies are conducted using different ecotype accessions, using different timepoints of seed development and using different tissue and RNA extraction methods (STable 4). Furthermore, genes not identified in this study may also have been excluded by filtering steps, such as absence of SNPs in accessions or biased accession expression. This is indeed a limitation for all studies restricted to experimental setups with few ecotype accessions. Excluding the genes that did not pass filtering steps in our setup, and therefore could not be analyzed, increased the overlap substantially (SFigure 6, SData 9). This suggests that experimental set-up and bioinformatic analysis considerably impact the resulting identified imprinted genes. Importantly, the largest overlap with our data is observed with studies that used similar endosperm tissue extraction methods (Del Toro-De León and Köhler, 2019; Picard et al., 2021).

### Overlap between temporal and spatial endosperm domains

As a next step, we compared the identities of imprinted genes recognized by our different FANS marker setups (EE, TE1, ESR) (Figure 4). This identified many genes that were uniquely called as imprinted in only one spatial or temporal domain (SData 10). On the other hand, only one MEG overlapped between EE, ESR and TE1 (SData 10). This gene, *SEEDSTICK (STK),* retains its imprinting state throughout seed development and ESR endosperm differentiation, and the relatively low number may suggest that imprinting is highly dynamic throughout seed development. Interestingly, STK, previously described as a transcription factor controlling ovule and seed integument identity (Ezquer et al., 2016), has recently also been identified to be imprinted in the endosperm by other FANS studies (Del Toro-De León and Köhler, 2019; Picard et al., 2021) and directly or indirectly regulates *BAN* and *TRANSPARENT TESTA 8 (TT8),* respectively (Mizzotti et al., 2014), both genes also identified as MEGs in this study.

**Figure 4.**
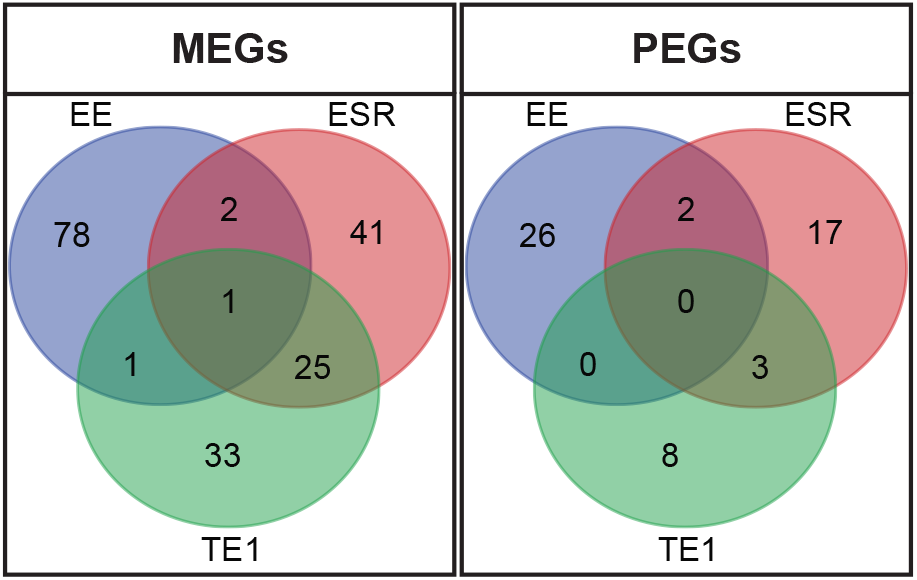
Overlapping MEGs and PEGs between EE, ESR and TE1. Venn diagrams display overlap between the identified imprinted genes in EE, ESR and TE1 experiments. Identified MEGs (left) and PEGs (right) between EE (blue), ESR (red) and TE1 (green) show temporal (EE and TE1), spatial (ESR and TE1) and spatio-temporal (EE, ESR and TE1) conservation of imprinting. Temporal (EE or TE1 specific) and spatial (ESR or TE1 specific) imprinting was observed for both MEGs and PEGs. Note the enhanced overlap between ESR and TE1.

Temporal differential imprinting dynamics is frequently observed in the comparison of allele specific profiles between early and late-stage markers (EE vs TE1 and EE vs ESR), where a low number of MEGs and PEGs overlap between early and late stages (four MEGs and two PEGs in both comparisons) (Figure 4, SData 10). In contrast, the allele specific profiles of the corresponding stage markers (TE1 vs ESR) were highly concordant, sharing 25 and three MEGs and PEGs, respectively (Figure 4). This suggests high accuracy of FANS since ESR represents a subdomain of the TE1 marker. Nevertheless, more than three times the amount of loci (74 MEGs and 25 PEGs) were imprinted only in the TE1 or in the ESR domains, suggesting spatial specific imprinting (SData 10). Although the overall expression levels, or lack of expression is not taken into account, both the temporal and spatial comparison suggest that genomic imprinting is highly dynamic, both during differentiation and development of the endosperm.

### Temporal dynamic regulation of genomic imprinting

In order to investigate if the parental specific allelic expression is truly regulated, and not just a consequence of stage specific repression or activation of genes, we next analyzed temporal regulation of imprinting by directly comparing the allelic expression values of imprinted genes between early and late developmental stages (EE vs TE1). To this end we required that a gene should be expressed in both FANS panels, but identified as imprinted (MEG or PEG) in one seed developmental stage whereas biparentally expressed (BEG) in the other seed developmental stage. Genes found to be imprinted at one time point and not expressed at the other time point were therefore not assessed in this analysis (SData 11).

The maternal:paternal fold change of (early) EE or (late) TE1 significantly imprinted genes (EE, TE1 and both EE and TE1; SData 11) were represented in a scatter plot (Figure 5). Three output scenarios can be proposed from this analysis; genes that are imprinted at both developmental time points, genes that are imprinted early but are biparentally expressed at the later developmental time-point and genes that are BEGs at the early developmental time point but imprinted at the later developmental time-point.

**Figure 5.**
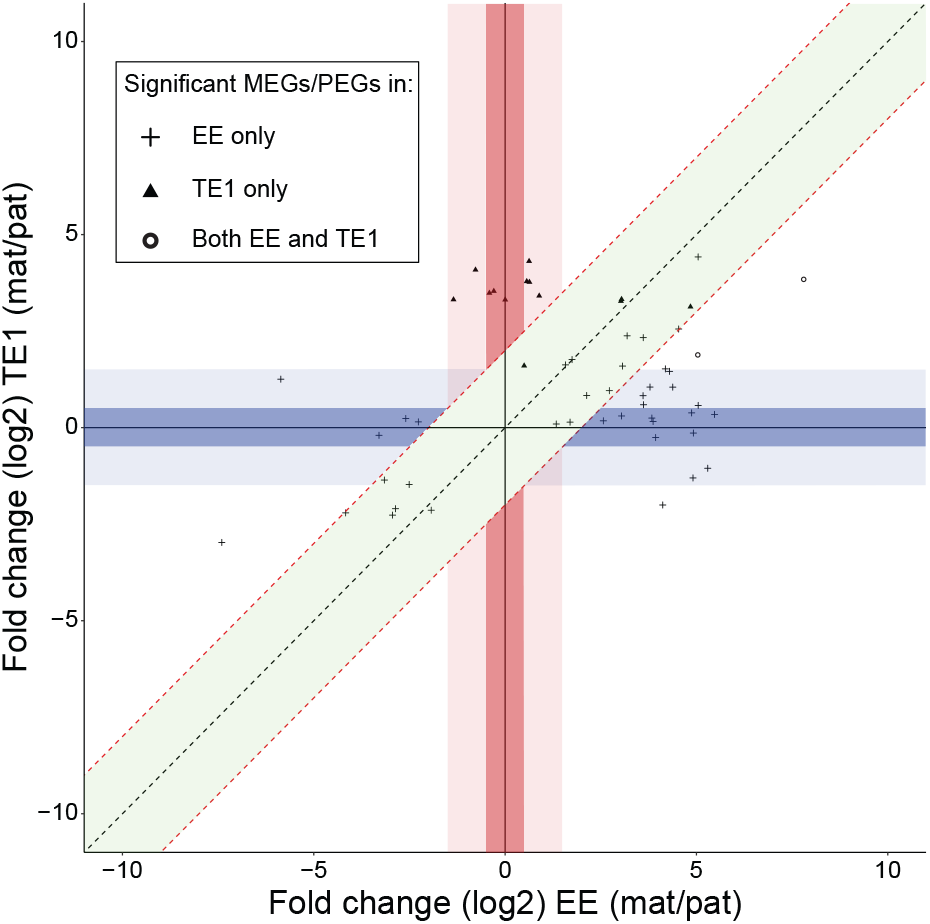
Identification of temporal regulation of imprinting. Scatter plot representation of the maternal:paternal fold change of early development (EE) or late development (TE1) significantly imprinted genes (EE, TE1 and both EE and TE1; SData 11). Genes called as MEGs or PEGs in one domain; EE (plus) or TE1 (triangle) or both domains EE and TE1 (circle) were included. Genes in the green area show a similar parental bias in both EE and TE1 (ΔFC (log2) < 2) indicating that they retain their imprinted state throughout seed development. Genes in the dark blue area are only imprinted in EE and are biparentally expressed in TE1 (0.5 < FC (log2) < 0.5). Genes in the light blue area are MEGs or PEGs for EE, but the parental bias in TE1 has not shifted to the same degree as for the dark blue area (−1.5 < FC (log2) < – 0.5 or 0.5 < FC (log2) < 1.5). Genes in the dark red area are only imprinted after cellurisation (TE1) and show biparental expression for EE (−0.5 < FC (log2) < 0.5). Concurrently, genes in the light red area are MEGs or PEGs for TE1, but parental bias in EE has not shifted to the same degree as for the dark red area (−1.5 < FC (log2) < −0.5 or 0.5 < FC (log2) < 1.5).

In our analysis, almost half of the genes (21/56) show a similar parental expression bias in EE and TE1 (Figure 5, diagonal green area), suggesting that they retain their imprinted state throughout seed development.

We identified eight MEGs and three PEGs, significantly imprinted in EE, that show biallelic expression in TE1 (Figure 5, horizontal dark blue area; Table 1A). Several of these EE specific MEGs have been previously identified as imprinted at a similar developmental stage (Del Toro-De León and Köhler, 2019; Picard et al., 2021), including *MYB7*, *BGAL1* and *TRM11* (SData 8). Most MEGs in EE show an increased expression in TE1 suggesting a reactivation of the paternal allele, presumably without any decrease in maternal expression (Table 1A). This is the case for all BEG turned MEGs in TE1 except for BGAL1 and AT3G58950 (SData 5). For the majority of these MEGs we thus infer that methylation marks are lost from paternal alleles, potentially gradually until later stages of seed development. Loss of methylation marks can be achieved by actively removing them exemplified by the 5mC DNA glycosylase family (Penterman et al., 2007). Another possibility is a discontinued maintenance of DNA methylation which will partly or completely remove methyl groups from a given allele depending on their cytosine context in the DNA. For EE PEGs, differential gene expression analysis between EE and TE1 (Table 1A, SData 2) revealed similar expression levels, and show that there is a simultaneous decrease of paternal and an increase of maternal expression (SData 5). Such a dynamic gene regulation scenario possibly involves several regulation mechanisms acting both through DNA demethylation, as described above, as well as through histone modifications. Histone modifications through the FIS-PRC2-complex are believed to mainly regulate the maternal allele of paternally expressed imprinted genes. Although only demonstrated for regulation in early in seed development it is possible that regulation of maternal alleles of PEGs are regulated by similar, or other, yet unknown mechanisms, early, and also at later seed developmental stages (Hornslien et al., 2019).

**Table 1.**
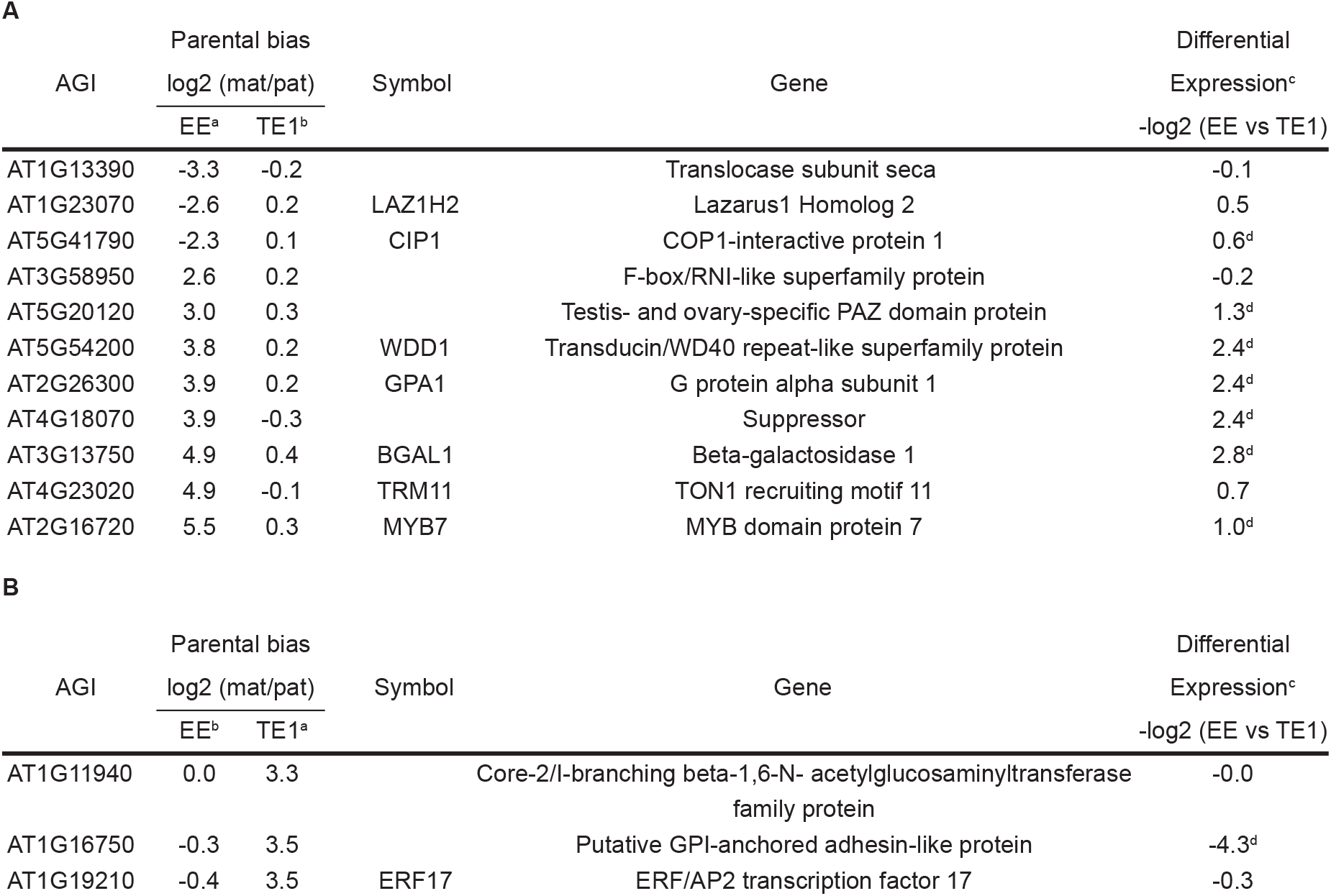
Temporal specific imprinted genes. Identified genes that show temporal dynamic regulation of genomic imprinting. A, Genes that are imprinted at endosperm development early stage (EE) and that are biparentally expressed at endosperm development late stage (TE1). B, Genes that are imprinted at endosperm development late stage (TE1) and that are biparentally expressed at endosperm development early stage (EE). ^a^ Significant imprinted MEGs and PEGs. FC (log2) between maternal and paternal informative reads within the domain. ^b^ Not significant. ^c^ Differential gene expression output between EE and TE1 total expression (SData 2) for selected genes. -FC (log2) values represent higher expression in EE (negative FC) or TE1 (positive FC). ^d^ Significant differentially expressed genes between EE and TE1.

Interestingly, three MEGs, significantly imprinted at the late developmental stage (TE1), displayed unbiased parental expression at the early endosperm developmental stage (EE) (Figure 5, vertical dark red area; Table 1B). Two out of these three MEGs show no significant change in overall expression level comparing EE and TE1 stages (Table 1B, SData 2). In both cases the paternal allele is completely silenced at the late time point whereas the maternal allele is constantly expressed (SData 5). These results suggest that the imprinting state is not predefined in the gametophyte but rather can arise at later seed developmental stages. Traditionally the paternal alleles of MEGs are thought to be silenced by DNA methylation, and in this scenario, *de novo* DNA methylation would be required to mediate silencing of the paternal allele in late seed development. The main methyltransferase shown to be involved in *de novo* methylation is DRM2 which indeed is highly upregulated (FC (log2) = 3.4) in our TE1 expression profile compared to our EE expression profile (SData 2), in line with previous endosperm expression studies (Belmonte et al., 2013).

### Spatial dynamic regulation of genomic imprinting

In contrast to the temporal, developmental stage comparison above (EE vs TE1), the comparison between imprinted loci from the same developmental stage but from partially overlapping endosperm domains (TE1 vs ESR) resulted in 29 imprinted loci (26 MEGs and three PEGs) with the same imprinting status in both domains (Figure 4). Nonetheless, spatial specific imprinting dominated, and a total of 104 genes (77 MEGs and 27 PEGs) were identified as imprinted in only TE1 or ESR domains (Figure 4, SData 10). We wanted to identify imprinted loci that displayed an actual bias in their parental contribution to the two domains investigated, and not merely were caused by the lack of expression in one of the domains. To this end, we performed a direct comparison of the allelic expression in the two domains. Genes that were not significantly expressed in both spatial domains compared were discarded (SData 12) and the remaining loci were required to be significantly identified as a MEG or PEG in one or both endosperm domains and at the same time display expression with no parental bias in the other domain.

The maternal:paternal fold change of ESR and TE1 significantly imprinted genes (ESR, TE1 and both ESR and TE1; SData 12) were represented in a scatter plot (Figure 6). Three types of spatial imprinting dynamics could be hypothesized from this analysis; loci that display the same imprinting status in the ESR and TE1 domains, loci that are MEGs or PEGs in TE1 and BEGs in ESR, and conversely MEGs or PEGs in ESR and BEGs in TE1.

**Figure 6.**
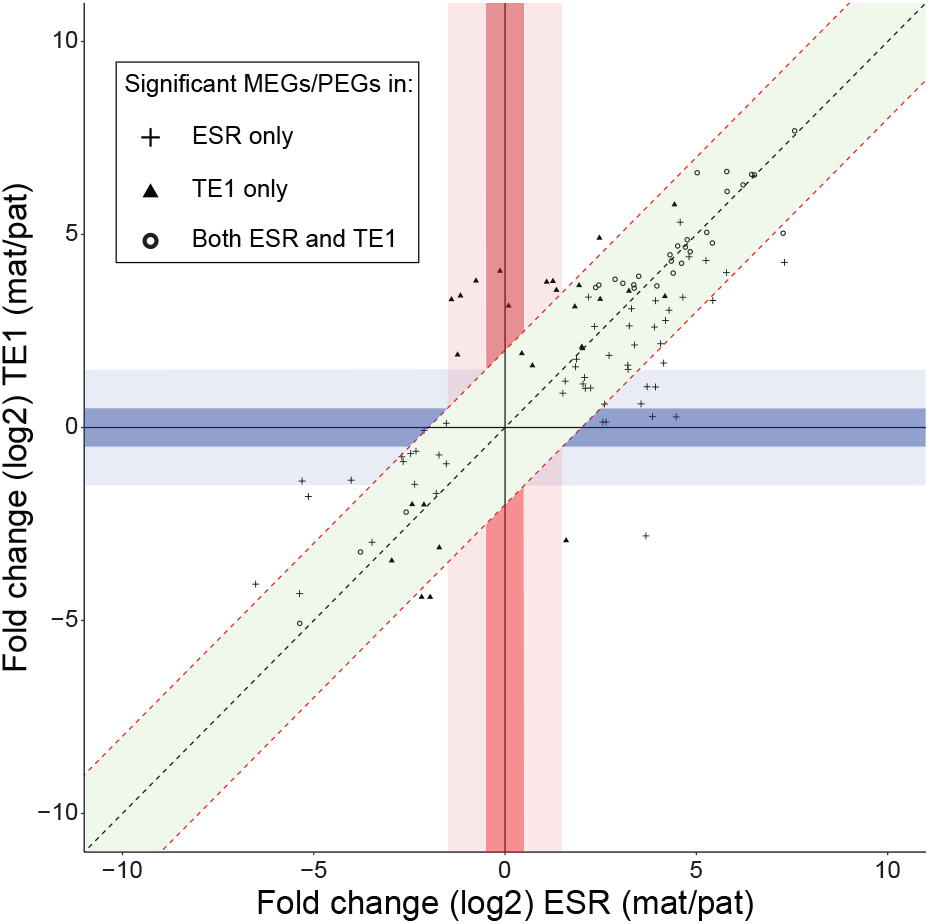
Identification of spatial specific imprinted genes. Scatter plot representation of the maternal:paternal fold change of total endosperm (TE1) and embryo surrounding region (ESR) significantly imprinted genes (in ESR, TE1 and both ESR and TE1; SData 12). Genes called as MEGs or PEGs in one domain; ESR (plus) or TE1 (triangle) or both domains ESR and TE1 (circle) were included. Genes in the green area show a similar parental bias in both ESR and TE1 (ΔFC (log2) < 2) indicating that their imprinting state is maintained throughout endosperm development reported by ESR and TE1. Genes in the dark blue area are imprinted in ESR but biparentally expressed in the overall total endosperm (TE1; −0.5 < FC (log2) < 0.5). Similarly, genes in the dark red area are imprinted in the overall total endosperm (TE1) but show biparental expression in the ESR (−0.5 < FC (log2) < 0.5). Genes in the light blue area are MEGs or PEGs for the ESR, but the parental bias in TE1 has not shifted to the same degree as for the dark blue area (−1.5 < FC (log2) < −0.5 or 0.5 < FC (log2) < 1.5). Concurrently, genes in the light red area are MEGs or PEGs for TE1, but parental bias in the ESR has not shifted to the same degree as for the dark red area (−1.5 < FC (log2) < −0.5 or 0.5 < FC (log2) < 1.5).

As expected due to the overlap between the domains, an absolute majority of genes (80/110) that are imprinted in either ESR, TE1 or both, and at the same time expressed in both domains show similar parental expression bias in ESR and TE1 (Figure 6, diagonal green area). 29 loci are significantly identified as imprinted in both the ESR and TE1 domain (Figure 6, circles in diagonal green area). The results also suggest that many of the loci called as significantly imprinted in only one of the two domains (Figure 4, SData 12) have a similar bias in the other domain, although not significant (Figure 6, triangle or plus in diagonal green area).

Interestingly, we identified five genes significantly imprinted in the ESR (four MEGs and one PEG), that are expressed in a biallelic manner in TE1 (Figure 6, horizontal dark blue area; Table 2A, SData 12). One of these loci, *AHL1,* has indeed previously been identified as imprinted in the endosperm (Del Toro-De León and Köhler, 2019; Hornslien et al., 2019) (SData 8). Albeit, these results may indicate that endosperm domain specific analysis, such as by the use of FANS, is imperative to successfully identify imprinted genes that otherwise would be masked by overall biparental expression in the endosperm. Differential expression analysis between ESR and TE1 showed that most loci were not significantly changed in overall expression (Table 2A, SData 2). For all genes identified as MEGs in ESR, the paternal allele is higher expressed in TE1 resulting in biparental expression in TE1, while the maternal allele of the identified MEGs are expressed at comparable levels in both TE1 and ESR (SData 5). Furthermore, we identified two significantly imprinted MEGs in TE1 that are expressed in a biallelic manner in ESR (Figure 6, horizontal dark red area; Table 2B, SData 12).

**Table 2.**
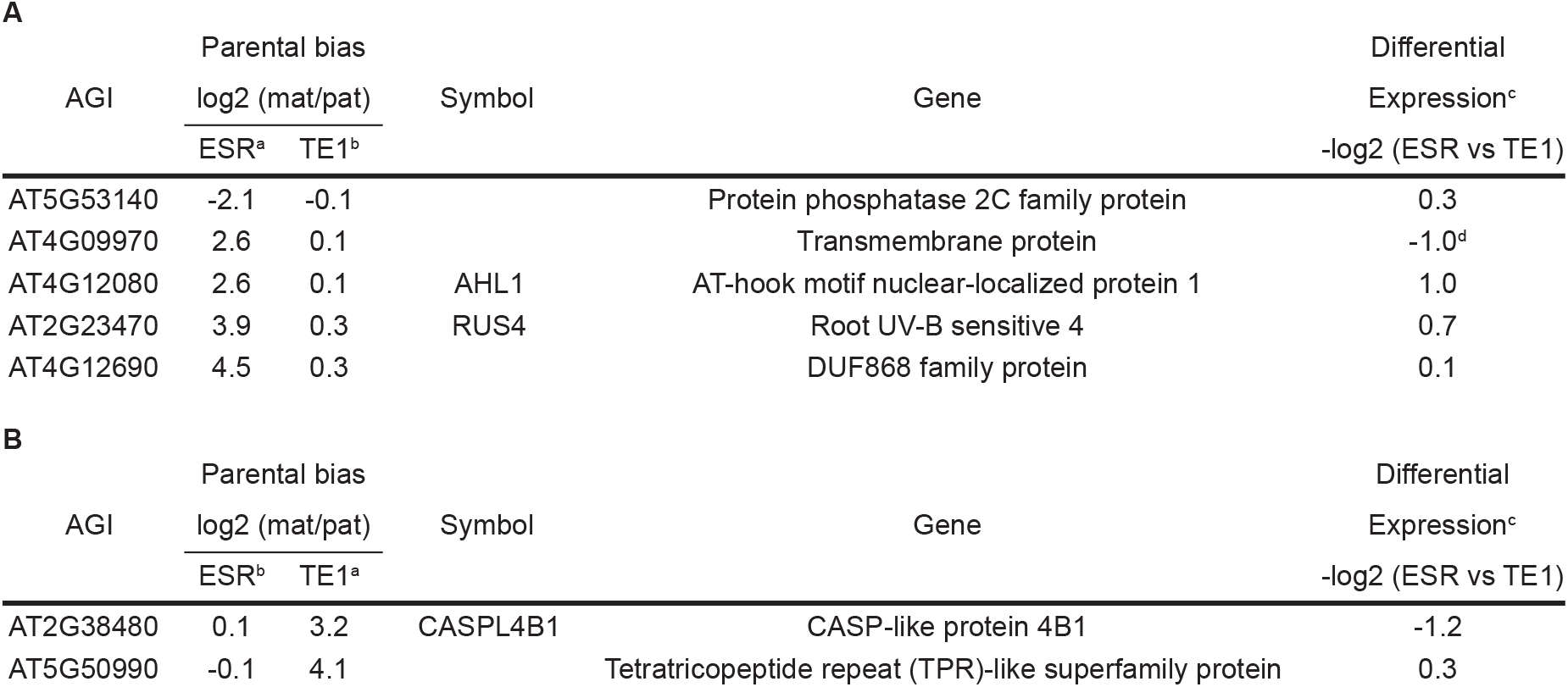
Spatial specific imprinted genes. Identified genes that show spatial dynamic regulation of genomic imprinting. A, Genes that are imprinted in the embryo surrounding region (ESR) and that are overall biparentally expressed in the total endosperm (TE1). B, Genes that are imprinted in the total endosperm (TE1) but that are biparentally expressed in the ESR. ^a^ Significant imprinted MEGs and PEGs. FC (log2) between maternal and paternal informative reads within the domain. ^b^ Not significant. ^c^ Differential gene expression output between ESR and TE1 total expression (SData 2) for selected genes. -FC (log2) values represent higher expression in ESR (negative FC) or TE1 (positive FC). ^d^ Significant differentially expressed genes between ESR and TE1.

| AGI | Parental bias |  | Symbol | Gene | Differential |
| --- | --- | --- | --- | --- | --- |
|  | log2 (mat/pat) |  |  |  | Expression<sup>c</sup> |
|  | ESR<sup>a</sup> | TE1<sup>b</sup> |  |  | -log2 (ESR vs TE1) |
| AT5G53140 | -2.1 | -0.1 |  | Protein phosphatase 2C family protein | 0.3 |
| AT4G09970 | 2.6 | 0.1 |  | Transmembrane protein | -1.0<sup>d</sup> |
| AT4G12080 | 2.6 | 0.1 | AHL1 | AT-hook motif nuclear-localized protein 1 | 1.0 |
| AT2G23470 | 3.9 | 0.3 | RUS4 | Root UV-B sensitive 4 | 0.7 |
| AT4G12690 | 4.5 | 0.3 |  | DUF868 family protein | 0.1 |

**Table 2. Spatial specific imprinted genes.**
| AGI | Parental bias |  | Symbol | Gene | Differential |
| --- | --- | --- | --- | --- | --- |
|  | log2 (mat/pat) |  |  |  | Expression<sup>c</sup> |
|  | ESR<sup>b</sup> | TE1<sup>a</sup> |  |  | -log2 (ESR vs TE1) |
| AT2G38480 | 0.1 | 3.2 | CASPL4B1 | CASP-like protein 4B1 | -1.2 |
| AT5G50990 | -0.1 | 4.1 |  | Tetratricopeptide repeat (TPR)-like superfamily protein | 0.3 |

Since the ESR is part of the TE1, the imprinting state of TE1 is affected by parental specific expression in the ESR. For genes that are imprinted in the ESR but identified as biallelic in the TE1 (Table 2A) we presume that the contribution of the ESR domain to the TE1 profile is minimal and that it will be masked by the TE1 contribution. Therefore, it is not unexpected that genes that are only imprinted in a sub domain of the endosperm are not detected as imprinted when looking at the endosperm as a whole. For genes that are imprinted in the total endosperm, but biallelic in the embryo surrounding region (Table 2B) we infer that the allelic bias must be established in a region of TE1 not including ESR. Indirectly, our results therefore indicate that this class of allelic expression identifies genes that are imprinted in a subdomain of TE1, excluding the ESR. Detection of imprinted genes in subregions of the endosperm as shown here, further indicate that the mechanisms underlying establishment, maintenance or release of imprinting may also act in a highly spatio-specific manner in the endosperm.

## Conclusion

Using FANS, we have successfully captured H2A-GFP domain specific nuclei from the seed, allowing the generation of pure endosperm expression profiles devoid of embryo and/or seed coat contamination which has been a challenge for studying imprinted genes in the endosperm. In addition, our data identifies loci that are spatially differentially imprinted in specific domains of endosperm. This suggests that regulation of genomic imprinting in the endosperm is more dynamic than previously reported and our findings are not readily explained by the canonical mechanisms for genomic imprinting.

In a systematic approach, genes have been assessed for their imprinting state at different seed developmental stages. We demonstrate temporal, developmental stage specific imprinting for several loci. Several genes were found to be imprinted in the syncytial endosperm while being biparentally expressed at a later seed developmental stage, as previously shown for *ZIX* (Ngo et al., 2012). This observation naturally follows the general assumption on how imprinting is established in gametes (Jullien et al., 2006; Gehring and Satyaki, 2017; Batista and Köhler, 2020), and that allelic bias is observed from early seed developmental stages. However, the systematic analysis provided here demonstrates that while many genes that are imprinted in early stages are completely silenced and switched off at later stages, some genes still retain a dynamic regulation by reactivation of the silenced allele or intricate balancing of the parental expression. Moreover, if expression of a gene remains at a constant level throughout endosperm development while the imprinting state changes, it is implied that both parental alleles are actively silenced and reactivated, suggesting that expression from the maternal and paternal allele is dynamically balanced.

Interestingly, we identified several genes to be biallelically expressed at early stages of seed development that acquire imprinted expression patterns through time. Most known mechanisms associated with genomic imprinting are primarily correlated to the establishment of imprinting marks in gametes, which do not explain this type of regulation. In order to accommodate such type of DNA methylation, a secondary mechanism able to distinguish parental alleles must be present. The RdDM pathway, resulting in small RNA directed *de novo* DNA methylation could be a potential regulatory mechanism for such a dynamic process (Kirkbride et al., 2019).

Furthermore, we show spatial dynamic regulation of genomic imprinting by providing direct and indirect evidence that some genes are imprinted only in a certain domain (ESR) of the endosperm or imprinted in a different endosperm domain than the ESR. Similar mechanisms as suggested above could be involved to reactivate the silenced allele in a domain specific manner. However, further investigation of this type of spatial imprinting is required to determine if the allele specific regulation is established early and then differentiate in distinct domains, or if the imprinting is established *de novo* after the specification of endosperm domain identities.

The future application of specific marker lines and FANS, as demonstrated in this report, allow for systematic dissection of endosperm subdomains in both a spatial and temporal manner. This will enhance our knowledge and understanding of discrete functions attributed to subdomains, and contribute to the discovery of novel imprinted genes that are not readily identified by investigating the complete endosperm. Ultimately, high-resolution analysis of domain specific parent-of-origin allelic expression may also contribute to resolve the evolutionary role of imprinting in the endosperm.

## Materials and Methods

### Plant material and growth conditions

Wild-type (WT) accessions Columbia (Col-0) and Tsushima (Tsu-1) seedlings were grown on MS-2 (Murashige and Skoog medium with 2% sucrose) plates in a 16-h-light / 8-h-dark cycle at 22°C for 10 days prior to transferring to soil. Plants were further grown on a 16-h-light / 8-h-dark cycle at 18°C. For the FANS experiment, Wild-type accession Tsu-1 and endosperm domain specific marker lines (Col-0) were grown on a 16-h-light / 8-h-dark cycle at 20-22°C.

### Identification and generation of endosperm marker lines

Endosperm domain specific marker lines were designed utilizing a two-component construct system where mGAL4-VP16 is expressed under control of a selected promoter sequence. The GAL4-VP16 transcription factor then activates expression of HISTONE 2A-GFP through the UAS regulatory element (Olvera-Carrillo et al., 2015). Candidate promoters were selected based on publicly available microarray data (Le et al., 2010) and on gene expression patterns (Winter et al., 2007). The promoters (STable 1) were obtained as Gateway cloning compatible amplicons from SAP collection (Benhamed et al., 2008) or amplified from genomic DNA using designated primers (STable 5). Promoter fragments were recombined into the pDONRP4P1r vector in a gateway BP reaction (Invitrogen). Subsequently, the promoter entry vectors were assembled together with the pENL1-GAL4-VP16-L2 entry vector into the pB-9FH2A-UAS-7m24GW destination vector in a multisite gateway LR reaction (Invitrogen), generating PROMOTER-OF-INTEREST:GAL4-VP16>>UAS:H2A-GFP constructs. The expression clones were transformed into *Agrobacterium tumefaciens* C58C1 (pMP90) competent cells using electroporation, which was used to transform wild-type Col-0 plants using the floral dip method (Clough and Bent, 1998). T1 plants were selected on antibiotics and segregation analysis was performed on T2 plants. All further analyses were performed with homozygous single-locus T3 plants of the following marker lines: Early Endosperm (EE; AT5G09370; proEE>>H2A-GFP); Embryo Surrounding Region (ESR; AT3G45130; proESR>>H2A-GFP); Developing Aleurone Layer (DAL; AT4G31060; proDAL>>H2A-GFP); Total Endosperm (TE1; AT4G00220; proTE1>>H2A-GFP).

### Imaging

For fluorescent microscopy, siliques of EE were manually dissected and seeds were imaged in water using a Zeiss Axioplan 2 imaging microscope equipped with a Zeiss Axio cam HDR camera. For confocal imaging of ESR, DAL and TE1 siliques were manually dissected and developing seeds were fixed in 4% paraformaldehyde (PFA), embedded in 5% agarose and 200 μm sections were obtained using a vibratome. Confocal images were acquired using an Axio Observer coupled to a LSM710 scanner with a Plan-Apochromat 20x/0.8 objective. GFP was excited with the 488 nm laser line of the Argon laser and the emission was detected between 495-545 nm. Autofluorescence in the seed was excited with the 633 nm and emission was detected between 638-721 nm. For staging of samples, light microscopy of developmental seed stages were performed using an Axioplan2 Imaging microscope equipped with a Zeiss Axio cam HDR camera. Wild-type Col-0 and Tsu-1 plants were emasculated two days prior to crossing with pollen from the same individual. Siliques were manually dissected at different timepoints using a stereomicroscope and seeds were mounted on a microscopy slide in a clearing solution of chloral hydrate in 30% glycerol as described previously (Grini et al., 2002).

### Fluorescence-Activated Nuclear Sorting (FANS)

Closed flower buds were emasculated two days prior to crossing with pollen from the paternal donor. Siliques were dissected at 4 days after pollination (4 DAP – EE) and 7 days after pollination (7 DAP – ESR; DAL; TE1) using a stereomicroscope and seeds were collected and chopped on ice in 50 μl Galbraith buffer (Galbraith et al., 1983) with 0.5% Triton X-100. Three different biological replicates were obtained with 40 siliques (EE; ESR) or 20 siliques (DAL; TE1) each. More Galbraith + Triton X-100 buffer (850 μl) was added before filtering through a 40 μm Partec filter. Propidium iodide (10 μg/μl) was added and GFP positive and negative nuclei were sorted directly into 500 μl RLT lysis buffer (Qiagen RNAeasy Plant Mini Kit) containing 1% 2-mercaptoethanol using a BD FACSAria™ III Cell Sorter (BD Biosciences).

### RNA isolation, library preparation and RNA sequencing of sorted nuclei

RNA was isolated as described in the RNeasy Plant Mini Kit (Qiagen) and 2 ng of total RNA was amplified with SMART-Seq v4 Ultra Low Input RNA kit (Clontech; version “091817”). Sequencing libraries were prepared with NEBnext Ultra DNA Library Prep Kit (New England Biolabs; version 6.0 – 2/18) using NEBNext Multiplex Oligos for Illumina-Dual Index Primers Set 1 (#E7600S) and libraries from three biological replicates (except for DAL-GFP+; two biological replicates) were equimolarly pooled and sequenced 150 bp paired end on a NovaSeq6000 S4 flow cell on one lane.

### Real-time Quantitative Polymerase Chain Reaction (RT-qPCR)

Nuclei from DAL were sorted for GFP positive (DAL GFP Pos) and GFP negative nuclei (DAL GFP Neg). Nuclei from ESR were sorted for GFP positive nuclei (ESR GFP Pos) and GFP negative nuclei, and the latter were re-sorted for 3C and 6C to capture only endosperm nuclei (ESR GFP Neg). RNA was extracted (Qiagen Micro RNeasy kit; Qiagen), cDNA was synthesized (iScript cDNA Synthesis Kit; BioRad) and RT-qPCR was performed for GFP with *UBIQUITIN10 (UBQ10)* as calibrator. The RT-qPCR was done in duplicate on a Lightcycler 480 (Roche) with SYBR green for detection (2,5 μl of mastermix, 0,25 μl of 5 μM of each forward and reverse primer, 1 μl H2O and 1 μl of cDNA). Data was analyzed with the -ΔΔCT method (Schmittgen and Livak, 2008), statistical analysis was done using the R package ‘dplyr’ (Wickham, 2018) and output was visualized in RStudio using the package ‘ggplot2’ (Wickham, 2016). The following primers were used for the RT-qPCR: GFP forward 5’ GACGGCAACTACAAGACCCG 3’; GFP reverse 5’ TTCAGCTCGATGCGGTTCAC 3’; UBQ10 forward 5’ TTCTGCCATCCTCCAACTGC 3’; UBQ10 reverse 5’ CACCCTCCACTTGGTCCTCA 3’.

### RNA isolation and sequencing for the generation of 4 DAP and 7 DAP reference transcriptomes of Col-0 and Tsu-1

Closed flower buds were emasculated two days prior to manual self-crossing. For the 4 DAP reference, siliques were dissected at 4 DAP for both wild-type accessions (Hornslien et al., 2019). For the 7 DAP reference, siliques were dissected at 7 and 8 DAP (Tsu-1) and 10 and 11 DAP (Col-0). Silique dissection was performed using a stereomicroscope and seeds were harvested in MagNA Lyser Green Beads tubes (Roche) tubes were collected into liquid nitrogen. Three different biological replicates were obtained from four different mother plants and twelve siliques per replicate. RNA was isolated as described in the Spectrum Total Plant RNA Kit (Sigma Aldrich) manual, Protocol A. During step 1, 1 ml Lysis Solution/2-mercaptoethanol solution was added to the tubes with the developing seeds. The tubes were shaken in a MagNA Lyser Instrument (Roche) at 7000 rounds per minute (rpm) for 15 sec, centrifuged at 13000 rpm in 4 °C for 15 sec and then placed at −20 °C for 2 minutes. This procedure was repeated three times before proceeding with the protocol. On-column DNase digestion was performed as described with On-Column DNaseI Digestion Set (Sigma Aldrich). For the 7 DAP reference, equal amounts of RNA from different timepoints were pooled for both accessions. RNA libraries were prepared using the Strand-Specific TruSeqTM RNA-Seq Library Prep (Illumina). Sequencing, 150 bp paired end, was performed over two lanes using an Illumina HiSeq 4000.

### Parent-of-origin specific differential gene expression analysis

A detailed description of bioinformatic analyses is provided as supplementary methods and scripts used in this study have been deposited to Github at https://github.com/PaulGrini/vanEkelenburg. In short, wild-type consensus reference transcriptomes for Col-0 and Tsu-1 were polished using pilon (Walker et al., 2014). Reads from heterozygous RNA endosperm domain marker lines were mapped to the generated reference transcriptomes with bowtie2 (Langmead and Salzberg, 2012). The Informative Read Pipeline (IRP) (Hornslien et al., 2019) was used to determine and extract informative reads. A read pair is informative, if it maps with indels to one reference transcriptome but without indels to the other reference transcriptome with at most one SNP. A read pair is also informative, if it maps without indels to both reference transcriptomes, but mapping to one of the transcriptomes results in fewer SNPs and the larger SNP count is more than twice the smaller count (i.e. 0 vs 1 SNP or 2 vs 5 SNPs). Statistical analysis to identify parent-of-origin specific differential gene expression bias was performed on informative reads with the R package limma (Ritchie et al., 2015). Bias was measured as fold change (log2) and significance was measured as p-value < 0.05 after adjustment for multiple tests

### Gene Ontology term enrichment analysis

Gene ontology (GO) term enrichment analysis was performed on the identified imprinted genes of the FANS markers using DAVID Bioinformatics Resources version 2021 (Huang et al., 2009a; Huang et al., 2009b) to identify significantly enriched GO terms using the EASE score (p-value < 0.05), a modified one-tailed Fisher’s exact t-test (Hosack et al., 2003).

### Image Analysis and Figure Preparation

Images were processed using Fiji (Schindelin et al., 2012). Figures were assembled in Adobe Illustrator 2021 (Adobe Systems Incorporated, San Jose, USA).

## Supplementary Figure legends

**SFigure 1. Fluorescence-activated nuclei sorting (FANS), ploidy analysis and marker identity verification.** Marker lines (EE, TE1, ESR and DAL) were analyzed for separation of GFP positive and negative nuclei. A, FANS separation profiles for EE, TE1, ESR and DAL. GFP positive and negative fractions are indicated in percent. B, FANS separation profiles for EE, TE1, ESR and DAL. GFP positive nuclei indicated in green and GFP negative fractions indicated in gray. C, Ploidy profiles of GFP positive (green) and GFP negative nuclei (grey). Succession of ploidy peaks (from left to right) are: diploid (gray), triploid (green), tetraploid (grey), hexaploid (green) and octoploid (grey). For ESR and DAL overlaps between multiples of diploid and triploid ploidies are expected since these markers do not capture total endosperm. D, RT-qPCR analysis of GFP transcripts in two replicates of ESR and DAL GFP positive nuclei (ESR GFP Pos and DAL GFP Pos respectively), and ESR GFP negative endosperm specific nuclei (ESR GFP Neg) and DAL GFP negative nuclei (DAL GFP Neg). UBQ10 was used as internal control and ESR GFP Pos and DAL GFP Pos were used as reference samples for ESR and DAL respectively. Error bars indicate standard deviation of the mean (SEM).

**SFigure 2. Spatio-temporal and seed developmental stage differential gene expression.** A, MA plot showing the fold change (log2) of differential expression between EE and ESR versus the mean expression (log2). The analysis revealed 8,714 genes to be significantly differentially expressed (red = up, blue = down, adjusted p-value < 0.05) within the spatiotemporal plane. B, MA plot showing the fold change (log2) differential expression between EE and TE1 versus the mean expression (log2). The analysis revealed 6,979 genes to be significantly differentially expressed (red = up, blue = down, adjusted p-value < 0.05) between seed developmental stages. Non significant (NS) genes are depicted in grey. Dashed lines indicate FC (log2) of 1 and −1.

**SFigure 3. Schematic overview of the applied informative read pipeline.** Sequenced reads from homozygous Col-0 and Tsu-1 and from heterozygous endosperm marker lines EE, TE1, ESR and DAL (blue), bioinformatic processing steps (red), filtering and normalization steps (brown) and final data output (green) are highlighted. 1) Homozygous reads were used to polish Col-0 and Tsu-1 reference transcriptomes for 4 and 7 DAP using pilon. 2) Homozygous reads from Col-0 and Tsu-1 were mapped to the reference transcriptomes with bowtie2 *-k 1* to identify significant gene expression between Col-0 and Tsu-1 using DESeq2. Genes that are not significantly expressed are retained. 3) Homozygous reads from Col-0 and Tsu-1 were mapped to the reference transcriptomes with bowtie2 *-k 2* to identify the genes for which informative reads preferentially map (>5 fold) to the correct parent. 4) Heterozygous reads from the marker lines were mapped to the reference transcriptomes with bowtie2 *-k 2* to call informative reads. 5) Heterozygous reads from the marker lines were mapped to the reference transcriptomes with bowtie2 *-k 2* to determine the library normalization factor for each replicate. 6) Heterozygous reads from the marker lines were mapped to the reference transcriptomes with bowtie2 *-k 1* to identify significant gene expression between the different endosperm domains using DESeq2. 7) Mitochondrial and chloroplast genes were removed and the homozygous gene filter and ecotype specific gene expression filter, generated in step 2 and 3 respectively, were applied. Informative read counts from the maternal allele (Col-0) were halved and all informative reads counts underwent library normalization, generated in step 5.

**SFigure 4. Ecotype specific gene expression analysis between Col-0 and Tsu-1 at different seed stages.** Differential expression between Col-0 and Tsu-1 at 4 DAP (A) and 7 DAP (B) to identify genes that are significantly ecotype preferentially expressed (adjusted p-value < 0.05). Non significant (NS) genes are depicted in grey. Dashed lines indicate fold change (log2) of 1 and −1.

**SFigure 5. Classification of genes based on fold change (FC) and statistical analysis.** Genes were subdivided as potential MEGs (left) or potential PEGs (right) if the informative read FC (log2) was > 0 or < 0 respectively. Genes with insufficient total informative read counts (≤30 including pseudocount), were classified as not enough informative reads. Genes were classified as Not biased if FC (log2) was < 1 or > −1 for MEGs and PEGs respectively. Genes were only considered as parentally biased if the informative read FC (log2) was ≥ 1 or ≤ −1 for MEGs and PEGs, respectively. If the adjusted p-value was not significant (≥ 0.05) genes were defined as Biased but not significant. Genes were determined to be imprinted if they showed a parental bias with FC (log2) ≥ 1 or ≤ −1 for MEGs and PEGs respectively and a significant adjusted p-value (< 0.05).

**SFigure 6. Overlap of identified imprinted genes with other studies increases after excluding genes that were not analyzed in this study.** MEGs (A) and PEGs (B) identified in EE, ESR, TE1 and all domains combined (All genes) were compared with imprinted gene lists from (Pignatta et al., 2014) (brown), (Hornslien et al., 2019) (red), (Del Toro-De León and Köhler, 2019) (green) and (Picard et al., 2021) (yellow). Analyzed genes = genes identified as imprinted in the other studies, but were not included in our analysis.

## Supplementary Table legends

**STable 1. Expression analysis of endosperm domain specific promoter marker line candidates.** Thirty-nine candidate promoter genes were selected based on microarray data (Le et al., 2010) and cloned in a dual component system where a histone2a-GFP fusion protein is expressed under control of an endosperm domain specific promoter. ^a^ GFP signal was observed throughout seed development and in each seed compartment. En = endosperm; Em = embryo; Sc = seed coat. ^b^ Seed stage at which GFP expression was observed. E = precellularization; L = post-cellularization; N/A = stage not specified. ^c^ GFP signal observed in different domains of the endosperm. EN = entire endosperm; ESR = embryo surrounding region; CH = chalazal endosperm; AL = aleurone layer; PEN = peripheral endosperm. ^d^ Constructs for which a GFP signal was observed in the embryo or seed coat are indicated with ‘x’. ^c/d^ Constructs for which no GFP signal was detected are indicated with ‘-’. ^e^ Marker name given to the promoter construct lines used for FANS. ESR = embryo surrounding region; TE1 = total endosperm; DAL = developing aleurone layer; EE = early endosperm. ^f^ A different set of long primers was used to amplify the promoter sequence. ^9^ GFP signal was observed in the entire endosperm, except the aleurone layer. ^h^ GFP was also observed in the embryo from bent cotyledon stage onwards.

**STable 2. Applying various filters reduced the final number of genes analyzed for parent-of-origin specific gene expression.** After calling informative reads, several filters were applied to remove genes that could result in false positives. Several mitochondrial and plastid genes, for which informative reads were detected, were removed. Genes for which reads from the same ecotype did not preferentially map to the correct parent (>5 fold) were excluded (homozygous mapping filter). Differentially expressed genes between whole seed Col-0 and Tsu-1 wild-type accessions were removed (ecotype filter). The remaining genes with informative reads were analyzed for parent-of-origin specific gene expression.

**STable 3. Gene Ontology (GO) term enrichment analysis**. GO term enrichment analysis performed with DAVID Bioinformatics Resources version 2021 (Huang et al., 2009a; Huang et al., 2009b). Significantly enriched GO terms were determined using the EASE score (p-value < 0.05; (Hosack et al., 2003)).

**STable 4. Summary of experimental set-up of various imprinting analysis studies.** ^a^ Weak MEGs and PEGs identified by (Picard et al., 2021) were not included. ^b^ Number indicates at which day after pollination tissue was collected and letter indicates developmental embryo stage (G: globular, LG: late globular, EH: early heart, LC: late cotyledon). ^c^ INTACT method to purify endosperm nuclei (Moreno-Romero et al., 2017). ^d^ Fluorescence-activated nuclear sorting of 3C/6C endosperm nuclei stained with Partec CyStain UV Precise P nuclei staining buffer. ^e^ Fluorescence-activated nuclear sorting of endosperm nuclei with GFP-tagged histones. ^f^ In addition, seed compartment transcript profiles were analyzed (Belmonte et al., 2013). ^9^ Tissue enrichment test (Schon and Nodine, 2017). ^h^ Ploidy analysis of 3C/6C specific endosperm nuclei peaks.

**STable 5. Cloning primers of the endosperm-domain marker construct lines.** Primers to clone promoter sequences for candidate promoter genes were designed or constructs were obtained from the SAP collection (Benhamed et al., 2008).

**STable 6. Library normalization factor for marker line replicates.** Directly after mapping with bowtie2, library normalization factors for each biological replicate of EE, ESR and TE1 were determined by dividing the total read number of each replicate by the average number of reads across all replicates.

## Notes

https://github.com/PaulGrini/vanEkelenburg

### Competing Interest Statement

The authors have declared no competing interest.

## References

Batista RA, Köhler C (2020) Genomic imprinting in plants—revisiting existing models. Genes Dev.

Belmonte MF, Kirkbride RC, Stone SL, Pelletier JM, Bui AQ, Yeung EC, Hashimoto M, Fei J, Harada CM, Munoz MD, et al (2013) Comprehensive developmental profiles of gene activity in regions and subregions of the Arabidopsis seed. Proc Natl Acad Sci U S A 110: E435–44

Benhamed M, Martin-Magniette M-L, Taconnat L, Bitton F, Servet C, De Clercq R, De Meyer B, Buysschaert C, Rombauts S, Villarroel R, et al (2008) Genome-scale Arabidopsis promoter array identifies targets of the histone acetyltransferase GCN5. Plant J 56: 493–504

Berger F, Grini PE, Schnittger A (2006) Endosperm: an integrator of seed growth and development. Curr Opin Plant Biol 9: 664–670

Birchler JA (1993) Dosage analysis of maize endosperm development. Annu Rev Genet 27: 181–204

Bjerkan KN, Hornslien KS, Johannessen IM, Krabberød AK, van Ekelenburg YS, Kalantarian M, Shirzadi R, Comai L, Brysting AK, Bramsiepe J, et al (2020) Genetic variation and temperature affects hybrid barriers during interspecific hybridization. Plant J 101:122-140

Brown RC, Lemmon BE, Nguyen H, Olsen O-A (1999) Development of endosperm in Arabidopsis thaliana. Sex Plant Reprod 12: 32–42

Clough SJ, Bent AF (1998) Floral dip: a simplified method forAgrobacterium-mediated transformation ofArabidopsis thaliana. Plant J 16: 735–743

Deal RB, Henikoff S (2010) The INTACT method for cell type-specific gene expression and chromatin profiling in Arabidopsis thaliana. Nat Protoc 6: 56–68

Del Toro-De León G, Köhler C (2019) Endosperm-specific transcriptome analysis by applying the INTACT system. Plant Reprod 32: 55–61

Ezquer I, Mizzotti C, Nguema-Ona E, Gotté M, Beauzamy L, Viana VE, Dubrulle N, Costa de Oliveira A, Caporali E, Koroney A-S, et al (2016) The Developmental Regulator SEEDSTICK Controls Structural and Mechanical Properties of the Arabidopsis Seed Coat. Plant Cell 28: 2478–2492

Feil R, Berger F (2007) Convergent evolution of genomic imprinting in plants and mammals. Trends Genet 23: 192–199

Galbraith DW, Harkins KR, Maddox JM, Ayres NM, Sharma DP, Firoozabady E (1983) Rapid flow cytometric analysis of the cell cycle in intact plant tissues. Science 220: 1049–1051

Gehring M, Satyaki PR (2017) Endosperm and Imprinting, Inextricably Linked. Plant Physiol 173: 143–154

Grini PE, Jürgens G, Hülskamp M (2002) Embryo and endosperm development is disrupted in the female gametophytic capulet mutants of Arabidopsis. Genetics 162: 1911–1925

Hornslien KS, Miller JR, Grini PE (2019) Regulation of Parent-of-Origin Allelic Expression in the Endosperm. Plant Physiol 180: 1498–1519

Hosack DA, Dennis G Jr, Sherman BT, Lane HC, Lempicki RA (2003) Identifying biological themes within lists of genes with EASE. Genome Biol 4: R70

Huang DW, Sherman BT, Lempicki RA (2009a) Systematic and integrative analysis of large gene lists using DAVID bioinformatics resources. Nat Protoc 4: 44–57

Huang DW, Sherman BT, Lempicki RA (2009b) Bioinformatics enrichment tools: paths toward the comprehensive functional analysis of large gene lists. Nucleic Acids Res 37: 1–13

Jacob Y, Mongkolsiriwatana C, Veley KM, Kim SY, Michaels SD (2007) The Nuclear Pore Protein AtTPR Is Required for RNA Homeostasis, Flowering Time, and Auxin Signaling. Plant Physiology 144: 1383–1390

Jullien PE, Kinoshita T, Ohad N, Berger F (2006) Maintenance of DNA methylation during the Arabidopsis life cycle is essential for parental imprinting. Plant Cell 18: 1360–1372

Kirkbride RC, Lu J, Zhang C, Mosher RA, Baulcombe DC, Chen ZJ (2019) Maternal small RNAs mediate spatial-temporal regulation of gene expression, imprinting, and seed development in Arabidopsis. Proc Natl Acad Sci U S A 116: 2761–2766

Langmead B, Salzberg SL (2012) Fast gapped-read alignment with Bowtie 2. Nat Methods 9: 357–359

Le BH, Cheng C, Bui AQ, Wagmaister JA, Henry KF, Pelletier J, Kwong L, Belmonte M, Kirkbride R, Horvath S, et al (2010) Global analysis of gene activity during Arabidopsis seed development and identification of seed-specific transcription factors. Proc Natl Acad Sci U S A 107: 8063–8070

Long Y, Liu Z, Jia J, Mo W, Fang L, Lu D, Liu B, Zhang H, Chen W, Zhai J (2021) FlsnRNA-seq: protoplasting-free full-length single-nucleus RNA profiling in plants. Genome Biol 22: 66

Mizzotti C, Ezquer I, Paolo D, Rueda-Romero P, Guerra RF, Battaglia R, Rogachev I, Aharoni A, Kater MM, Caporali E, et al (2014) SEEDSTICK is a Master Regulator of Development and Metabolism in the Arabidopsis Seed Coat. PLoS Genetics 10: e1004856

Moreno-Romero J, Santos-González J, Hennig L, Köhler C (2017) Applying the INTACT method to purify endosperm nuclei and to generate parental-specific epigenome profiles. Nat Protoc 12: 238–254

Ngo QA, Baroux C, Guthörl D, Mozerov P, Collinge MA, Sundaresan V, Grossniklaus U (2012) The Armadillo repeat gene ZAK IXIK promotes Arabidopsis early embryo and endosperm development through a distinctive gametophytic maternal effect. Plant Cell 24: 4026–4043

Nowack MK, Ungru A, Bjerkan KN, Grini PE, Schnittger A (2010) Reproductive crosstalk: seed development in flowering plants. Biochem Soc Trans 38: 604–612

Olvera-Carrillo Y, Van Bel M, Van Hautegem T, Fendrych M, Huysmans M, Simaskova M, van Durme M, Buscaill P, Rivas S, Coll NS, et al (2015) A Conserved Core of Programmed Cell Death Indicator Genes Discriminates Developmentally and Environmentally Induced Programmed Cell Death in Plants. Plant Physiol 169: 2684–2699

Penterman J, Zilberman D, Huh JH, Ballinger T, Henikoff S, Fischer RL (2007) DNA demethylation in the Arabidopsis genome. Proc Natl Acad Sci U S A 104: 6752–6757

Picard CL, Povilus RA, Williams BP, Gehring M (2021) Transcriptional and imprinting complexity in Arabidopsis seeds at single-nucleus resolution. Nat Plants 7: 730–738

Pignatta D, Erdmann RM, Scheer E, Picard CL, Bell GW, Gehring M (2014) Natural epigenetic polymorphisms lead to intraspecific variation in Arabidopsis gene imprinting. Elife 3: e03198

Ritchie ME, Phipson B, Wu D, Hu Y, Law CW, Shi W, Smyth GK (2015) limma powers differential expression analyses for RNA-sequencing and microarray studies. Nucleic Acids Res 43: e47

Satyaki PRV, Gehring M (2019) Paternally Acting Canonical RNA-Directed DNA Methylation Pathway Genes Sensitize Arabidopsis Endosperm to Paternal Genome Dosage. Plant Cell 31: 1563–1578

Satyaki PRV, Gehring M (2017) DNA methylation and imprinting in plants: machinery and mechanisms. Crit Rev Biochem Mol Biol 52: 163–175

Schindelin J, Arganda-Carreras I, Frise E, Kaynig V, Longair M, Pietzsch T, Preibisch S, Rueden C, Saalfeld S, Schmid B, et al (2012) Fiji: an open-source platform for biological-image analysis. Nat Methods 9: 676–682

Schmittgen TD, Livak KJ (2008) Analyzing real-time PCR data by the comparative C(T) method. Nat Protoc 3: 1101–1108

Schon MA, Nodine MD (2017) Widespread Contamination of Arabidopsis Embryo and Endosperm Transcriptome Data Sets. Plant Cell 29: 608–617

Shirzadi R, Andersen ED, Bjerkan KN, Gloeckle BM, Heese M, Ungru A, Winge P, Koncz C, Aalen RB, Schnittger A, et al (2011) Genome-Wide Transcript Profiling of Endosperm without Paternal Contribution Identifies Parent-of-Origin-Dependent Regulation of AGAMOUS-LIKE36. PLoS Genet 7: e1001303

Vu TM, Nakamura M, Calarco JP, Susaki D, Lim PQ, Kinoshita T, Higashiyama T, Martienssen RA, Berger F (2013) RNA-directed DNA methylation regulates parental genomic imprinting at several loci in Arabidopsis. Development 140: 2953–2960

Walker BJ, Abeel T, Shea T, Priest M, Abouelliel A, Sakthikumar S, Cuomo CA, Zeng Q, Wortman J, Young SK, et al (2014) Pilon: an integrated tool for comprehensive microbial variant detection and genome assembly improvement. PLoS One 9: e112963

Weijers D, Van Hamburg J-P, Van Rijn E, Hooykaas PJJ, Offringa R (2003) Diphtheria toxin-mediated cell ablation reveals interregional communication during Arabidopsis seed development. Plant Physiol 133: 1882–1892

Wickham H (2018) dplyr: A Grammar of Data Manipulation. R package version 0.7.6. https://CRAN.R-project.org/package=dplyr

Wickham H (2016) ggplot2: Elegant Graphics for Data Analysis. Springer-Verlag New York

Winter D, Vinegar B, Nahal H, Ammar R, Wilson GV, Provart NJ (2007) An “Electronic Fluorescent Pictograph” Browser for Exploring and Analyzing Large-Scale Biological Data Sets. PLoS One 2: e718

Wolff P, Jiang H, Wang G, Santos-González J, Köhler C (2015) Paternally expressed imprinted genes establish postzygotic hybridization barriers in Arabidopsis thaliana. Elife. doi: 10.7554/eLife.10074

Zhang S, Wang D, Zhang H, Skaggs MI, Lloyd A, Ran D, An L, Schumaker KS, Drews GN, Yadegari R (2018) FERTILIZATION-INDEPENDENT SEED-Polycomb Repressive Complex 2 Plays a Dual Role in Regulating Type I MADS-Box Genes in Early Endosperm Development. Plant Physiol 177: 285–299

