## Supplementary Figures for "Spatial and Temporal Regulation of Parent-of-Origin Allelic Expression in the Endosperm"

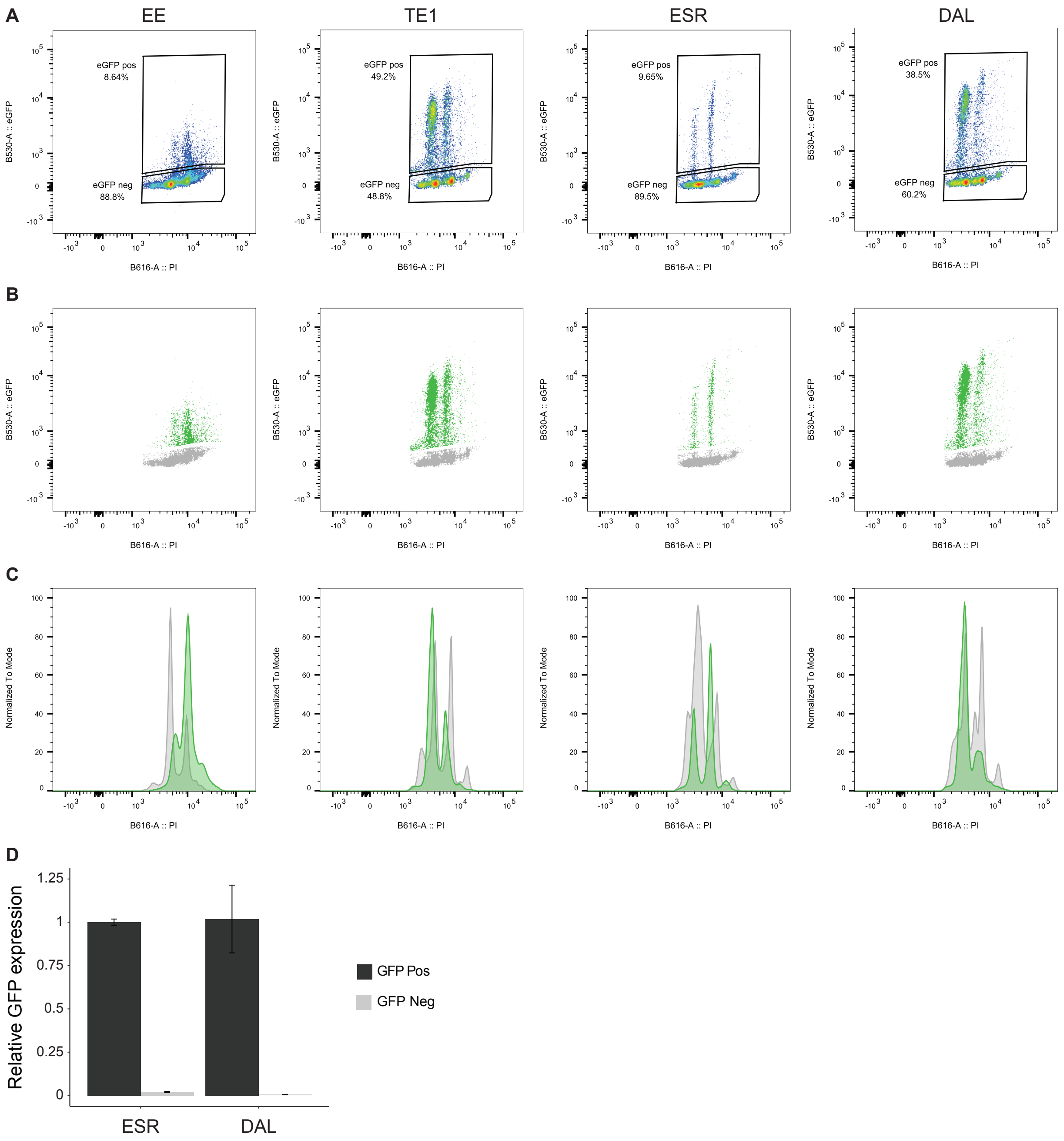


**SFigure 1. Fluorescence-activated nuclei sorting (FANS), ploidy analysis and marker identity verification.** Marker lines (EE, TE1, ESR and DAL) were analyzed for separation of GFP positive and negative nuclei. A, FANS separation profiles for EE, TE1, ESR and DAL. GFP positive and negative fractions are indicated in percent. B, FANS separation profiles for EE, TE1, ESR and DAL. GFP positive nuclei indicated in green and GFP negative fractions indicated in gray. C, Ploidy profiles of GFP positive (green) and GFP negative nuclei (grey). Succession of ploidy peaks (from left to right) are: diploid (gray), triploid (green), tetraploid (grey), hexaploid (green) and octoploid (grey). For ESR and DAL overlaps between multiples of diploid and triploid ploidies are expected since these markers do not capture total endosperm. D, RT-qPCR analysis of GFP transcripts in two replicates of ESR and DAL GFP positive nuclei (ESR GFP Pos and DAL GFP Pos respectively), and ESR GFP negative endosperm specific nuclei (ESR GFP Neg) and DAL GFP negative nuclei (DAL GFP Neg). UBQ10 was used as internal control and ESR GFP Pos and DAL GFP Pos were used as reference samples for ESR and DAL respectively. Error bars indicate standard deviation of the mean (SEM).

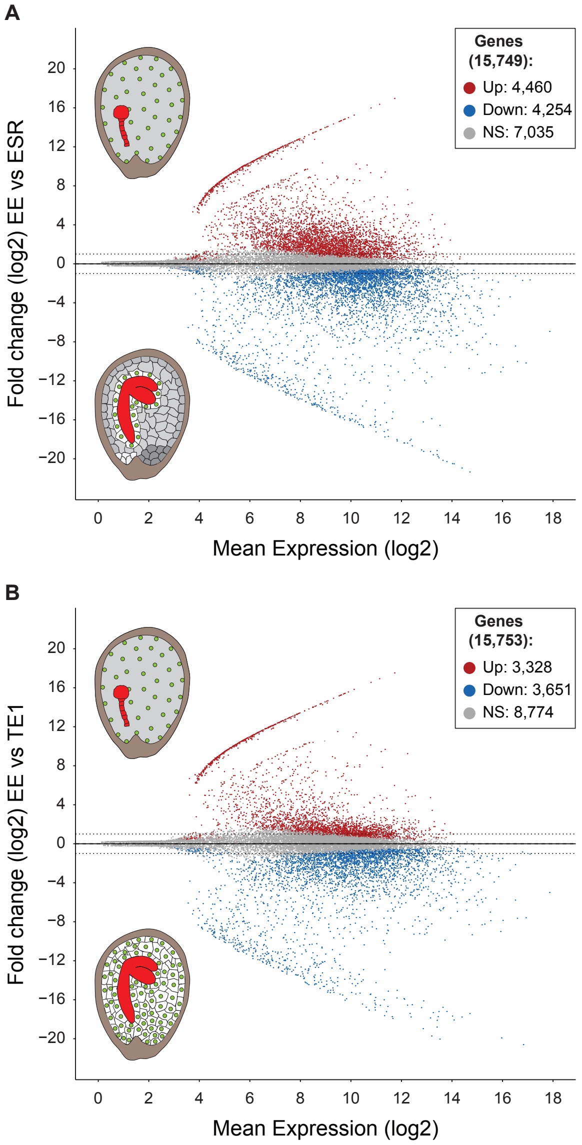


**SFigure 2. Spatio-temporal and seed developmental stage differential gene expression.** A, MA plot showing the fold change (log2) of differential expression between EE and ESR versus the mean expression (log2). The analysis revealed 8,714 genes to be significantly differentially expressed (red = up, blue = down, adjusted p-value < 0.05) within the spatio-temporal plane. B, MA plot showing the fold change (log2) differential expression between EE and TE1 versus the mean expression (log2). The analysis revealed 6,979 genes to be significantly differentially expressed (red = up, blue = down, adjusted p-value < 0.05) between seed developmental stages. Non significant (NS) genes are depicted in grey. Dashed lines indicate FC (log2) of 1 and -1.

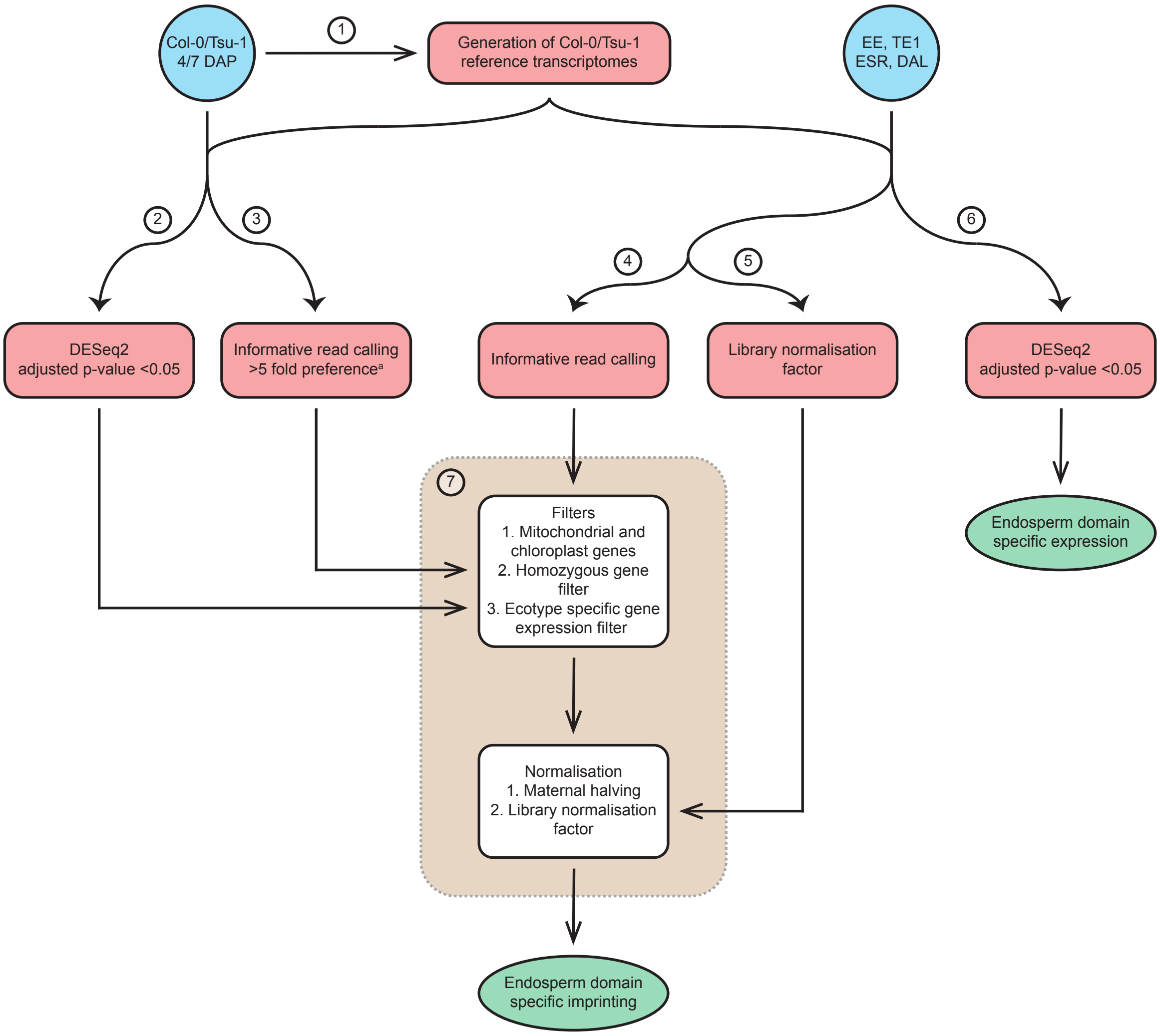


**SFigure 3. Schematic overview of the applied informative read pipeline.** Sequenced reads from homozygous Col-0 and Tsu-1 and from heterozygous endosperm marker lines EE, TE1, ESR and DAL (blue), bioinformatic processing steps (red), filtering and normalization steps (brown) and final data output (green) are highlighted. 1) Homozygous reads were used to generate Col-0 and Tsu-1 reference transcriptomes for 4 and 7 DAP using pilon. 2) Homozygous reads from Col-0 and Tsu-1 were mapped to the reference transcriptomes with bowtie2 *-k 1* to identify significant gene expression between Col-0 and Tsu-1 using DESeq2. Genes that are not significantly expressed are retained. 3) Homozygous reads from Col-0 and Tsu-1 were mapped to the reference transcriptomes with bowtie2 *-k 2* to identify the genes for which informative reads preferentially map (>5 fold) to the correct parent. 4) Heterozygous reads from the marker lines were mapped to the reference transcriptomes with bowtie2 *-k 2* to call informative reads. 5) Heterozygous reads from the marker lines were mapped to the reference transcriptomes with bowtie2 *-k 2* to determine the library normalization factor for each replicate. 6) Heterozygous reads from the marker lines were mapped to the reference transcriptomes with bowtie2 *-k 1* to identify significant gene expression between the different endosperm domains using DESeq2. 7) Mitochondrial and chloroplast genes were removed and the homozygous gene filter and ecotype specific gene expression filter, generated in step 2 and 3 respectively, were applied. Informative read counts from the maternal allele (Col-0) were halved and all informative reads counts underwent library normalization, generated in step 5.

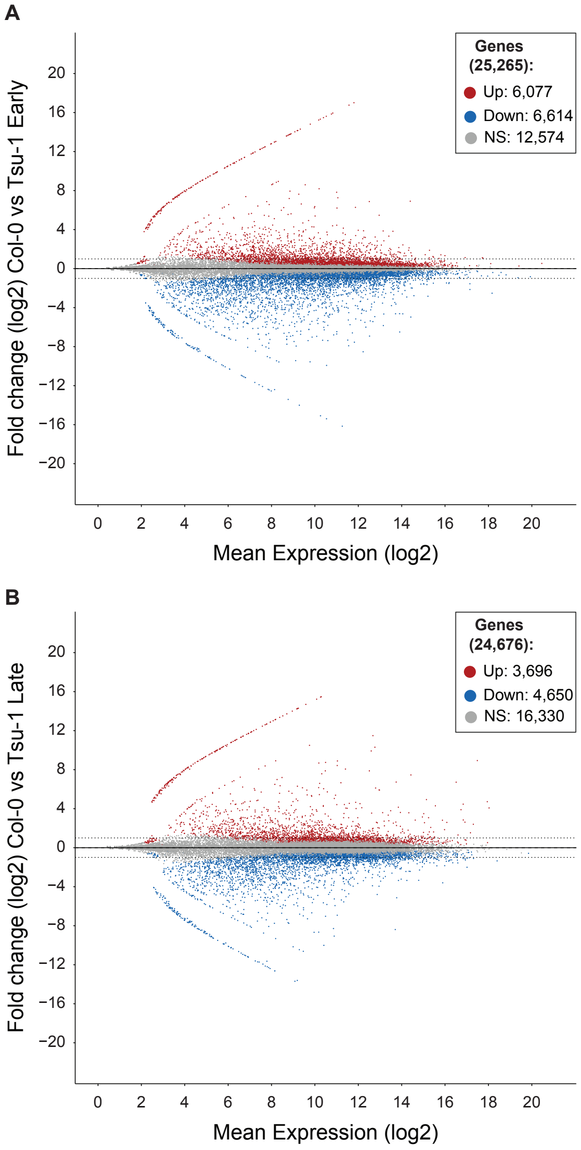


**SFigure 4. Ecotype specific gene expression analysis between Col-0 and Tsu-1 at different seed stages.** Differential expression between Col-0 and Tsu-1 at 4 DAP (A) and 7 DAP (B) to identify genes that are significantly ecotype preferentially expressed (adjusted p-value $< 0.05$). Non significant (NS) genes are depicted in grey. Dashed lines indicate fold change (log2) of 1 and -1.

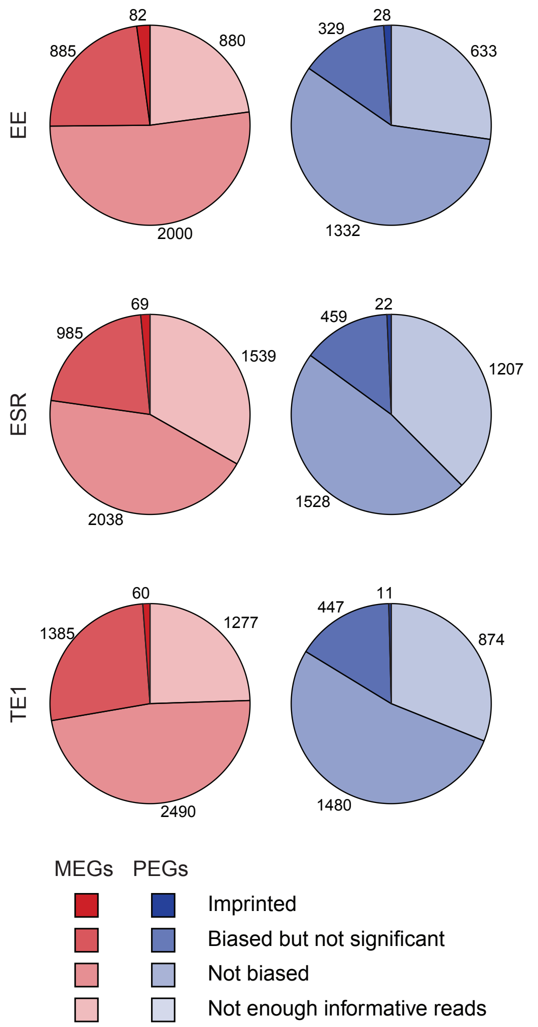


**SFigure 5. Classification of genes based on fold change (FC) and statistical analysis.** Genes were subdivided as potential MEGs (left) or potential PEGs (right) if the informative read FC (log2) was > 0 or < 0 respectively. Genes with insufficient total informative read counts (≤30 including pseudocount), were classified as not enough informative reads. Genes were classified as Not biased if FC (log2) was < 1 or > -1 for MEGs and PEGs respectively. Genes were only considered as parentally biased if the informative read FC (log2) was ≥ 1 or ≤ -1 for MEGs and PEGs, respectively. If the adjusted p-value was not significant (≥ 0.05) genes were defined as Biased but not significant. Genes were determined to be imprinted if they showed a parental bias with FC (log2) ≥ 1 or ≤ -1 for MEGs and PEGs respectively and a significant adjusted p-value (< 0.05).

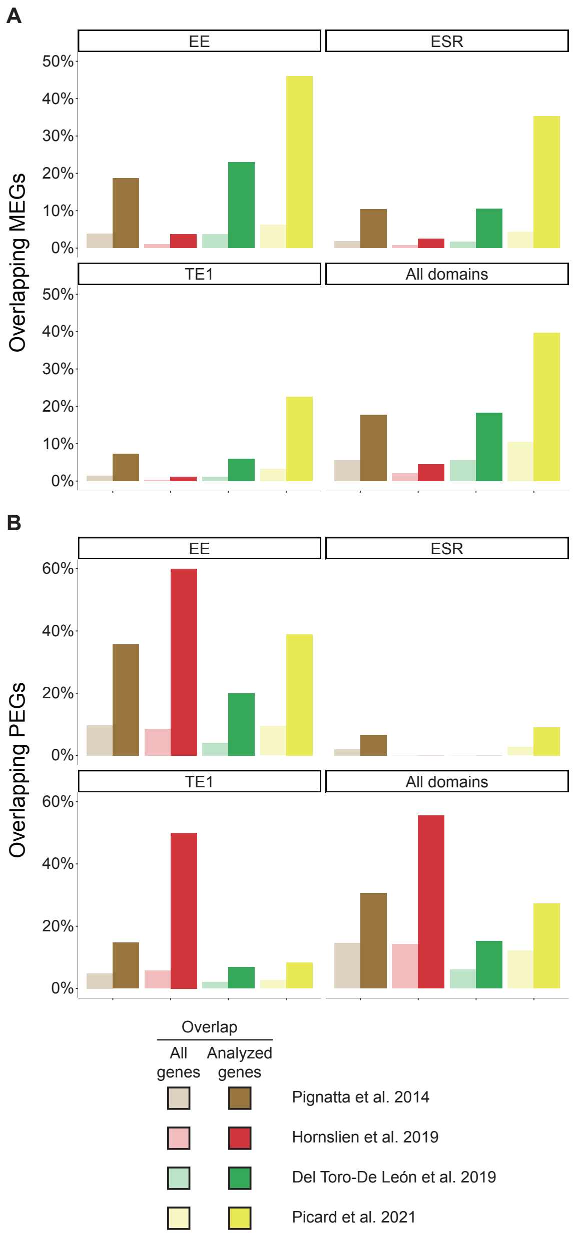


**SFigure 6. Overlap of identified imprinted genes with other studies increases after excluding genes that were not analyzed in this study.** MEGs (A) and PEGs (B) identified in EE, ESR, TE1 and all domains combined (All genes) were compared with imprinted gene lists from (Pignatta et al., 2014) (brown), (Hornslien et al., 2019) (red), (Del Toro-De León and Köhler, 2019) (green) and (Picard et al., 2021) (yellow). Analyzed genes = genes identified as imprinted in the other studies, but were not included in our analysis.
