## Supplementary Tables for "Spatial and Temporal Regulation of Parent-of-Origin Allelic Expression in the Endosperm"

| AGI | Symbol | Gene name | Expression[a] | | | | Marker[e] |
|---|---|---|---|---|---|---|---|
| | | | Stage[b] | En[c] | Em[d] | Sc[d] | |
| AT1G02450 | NIMIN1 | NIM1-INTERACTING 1 | E | EN | - | - | |
| AT1G11190 | BFN1 | BIFUNCTIONAL NUCLEASE I | N/A | ESR | - | - | |
| AT1G17020 | SRG1 | SENESCENCE-RELATED GENE 1 | L | ESR; CH; AL | - | - | |
| AT1G25410 | IPT6 | ISOPENTENYLTRANSFERASE 6 | L | AL | x | - | |
| AT1G48930 | GH9C1 | GLYCOSYL HYDROLASE 9C1 | L | EN | - | - | |
| AT1G62080 | TBA3 | TESTA ABUNDANT3 | L | - | - | x | |
| AT1G68510 | LBD42 | LOB DOMAIN-CONTAINING PROTEIN 42 | E | PEN | x | - | |
| AT2G25940 | ALPHA-VPE | ALPHA-VACUOLAR PROCESSING ENZYME | L | AL | - | - | |
| AT2G27380 | EPR1 | EXTENSIN PROLINE-RICH 1 | N/A | EN | - | - | |
| AT2G34315 | | Avirulence induced family protein | N/A | - | - | - | |
| AT2G38900 | | Predicted to encode a pathogenesis-related peptide that belongs to the PR-6 proteinase inhibitor family | N/A | - | - | x | |
| AT2G41290 | SSL2 | STRICTOSIDINE SYNTHASE-LIKE 2 | N/A | EN | x | x | |
| AT2G42430 | LBD16 | LATERAL ORGAN BOUNDARIES-DOMAIN 16 | L | AL | - | - | |
| AT2G43660 | | Carbohydrate-binding X8 domain superfamily protein | E | EN | - | - | |
| AT3G12110 | ACT11 | ACTIN-11 | N/A | EN | x | - | |
| AT3G19660 | | Hypothetical protein | L | AL | - | - | |
| AT3G22540 | | Hypothetical protein (DUF1677) | L | ESR | x | - | |
| AT3G29190 | | Terpenoid cyclases/Protein prenyltransferases superfamily protein | L | AL | - | - | |
| AT3G29770 | MES11 | METHYL ESTERASE 11 | L | EN | x | - | |
| AT3G45130 | LAS1 | LANOSTEROL SYNTHASE 1 | L | ESR | - | - | ESR |
| AT3G57540 | REM4.1 | REMORIN GROUP 4 1 | L | AL | x | - | |
| AT3G60245 | | Zinc-binding ribosomal protein family protein | N/A | EN | x | - | |
| AT4G00180 | YAB3 | YABBY3 | N/A | - | x | - | |
| AT4G00220 | JLO | JAGGED LATERAL ORGANS | L | EN | - | - | TE1 |
| AT4G04460 | PASPA3 | PUTATIVE ASPARTIC PROTEINASE A3 | L | EN | - | - | |
| AT4G05320 | UBQ10 | POLYUBIQUITIN 10 | N/A | EN | x | x | |
| AT4G21980 | APG8A | AUTOPHAGY 8A | N/A | - | - | x | |
| AT4G30960 | SIP3 | SOS3-INTERACTING PROTEIN 3 | N/A | EN | - | - | |
| AT4G30970 | | Hypothetical protein | N/A | EN | x | - | |
| AT4G30970[f] | | Hypothetical protein | N/A | - | x | - | |
| AT4G31060 | | Member of the DREB subfamily A-5 of ERF/AP2 transcription factor family | L | AL | - | - | DAL |
| AT5G09370 | LTPG29 | GLYCOSYLPHOSPHATIDYLINOSITOL-ANCHORED LIPID PROTEIN TRANSFER 29 | E,L | EN | - | - | EE |
| AT5G10220 | ANN6 | ANNEXIN 6 | E,L | EN[g] | - | - | |
| AT5G13600 | | Phototropic-responsive NPH3 family protein | E,L | EN | x[h] | - | |
| AT5G45890 | SAG12 | SENESCENCE-ASSOCIATED GENE 12 | E,L | - | - | x | |
| AT5G46950 | INVINH2 | INVERTASE INHIBITOR 2 | E,L | EN | x | - | |
| AT5G50260 | CEP1 | CYSTEINE ENDOPEPTIDASE 1 | N/A | EN | x | x | |
| AT5G54650 | FH5 | FORMIN HOMOLOGY 5 | E | EN | - | - | |
| AT5G56620 | NAC099 | NAC DOMAIN CONTAINING PROTEIN 99 | N/A | CH | - | - | |

**STable 1. Expression analysis of endosperm domain specific promoter marker line candidates.** Thirty-nine candidate promoter genes were selected based on microarray data (Le et al., 2010) and cloned in a dual component system where a histone2a-GFP fusion protein is expressed under control of an endosperm domain specific promoter. [a] GFP signal was observed throughout seed development and in each seed compartment. En = endosperm; Em = embryo; Sc = seed coat. [b] Seed stage at which GFP expression was observed. E = precellularization; L = post-cellularization; N/A = stage not specified. [c] GFP signal observed in different domains of the endosperm. EN = entire endosperm; ESR = embryo surrounding region; CH = chalazal endosperm; AL = aleurone layer; PEN = peripheral endosperm. [d] Constructs for which a GFP signal was observed in the embryo or seed coat are indicated with ‘x’. c/d Constructs for which no GFP signal was detected are indicated with ‘-’. [e] Marker name given to the promoter construct lines used for FANS. ESR = embryo surrounding region; TE1 = total endosperm; DAL = developing aleurone layer; EE = early endosperm. [f] A different set of long primers was used to amplify the promoter sequence. [g] GFP signal was observed in the entire endosperm, except the aleurone layer. [h] GFP was also observed in the embryo from bent cotyledon stage onwards.

| Marker line | # Genes with informative reads | # Genes excluded after filter applied | | | | # Genes analyzed |
|---|---|---|---|---|---|---|
| | | Mitochondrial | Plastid | Homozygous mapping filter | Ecotype filter | |
| EE | 14,962 | 27 | 19 | 932 | 7,815 | 6,169 |
| ESR | 13,510 | 17 | 17 | 677 | 4,952 | 7,847 |
| TE1 | 13,817 | 17 | 19 | 700 | 5,057 | 8,024 |

**STable 2. Applying various filters reduced the final number of genes analyzed for parent-of-origin specific gene expression.** After calling informative reads, several filters were applied to remove genes that could result in false positives. Several mitochondrial and plastid genes, for which informative reads were detected, were removed. Genes for which reads from the same ecotype did not preferentially map to the correct parent (>5 fold) were excluded (homozygous mapping filter). Differentially expressed genes between whole seed Col-0 and Tsu-1 wild-type accessions were removed (ecotype filter). The remaining genes with informative reads were analyzed for parent-of-origin specific gene expression.

| Domain | MEGs/PEGs | GO molecular function | Fold enrichment | p-value |
|---|---|---|---|---|
| EE | MEGs | RNA polymerase II core promoter proximal region sequence-specific DNA binding | 6.1 | 8.3E-03 |
| EE | MEGs | RNA polymerase II transcription factor activity, sequence-specific DNA binding | 5.5 | 1.2E-02 |
| EE | MEGs | transcription factor activity, sequence-specific DNA binding | 2.5 | 7.3E-03 |
| EE | PEGs | poly(A) binding | 111.3 | 1.7E-02 |
| EE | MEGs + PEGs | RNA polymerase II core promoter proximal region sequence-specific DNA binding | 4.9 | 1.9E-02 |
| EE | MEGs + PEGs | RNA polymerase II transcription factor activity, sequence-specific DNA binding | 4.3 | 2.7E-02 |
| EE | MEGs + PEGs | transcription factor activity, sequence-specific DNA binding | 1.9 | 3.9E-02 |
| ESR | MEGs | sequence-specific DNA binding | 5.1 | 1.5E-02 |
| ESR | MEGs + PEGs | sequence-specific DNA binding | 3.7 | 4.2E-02 |
| TE1 | MEGs | transcription factor activity, sequence-specific DNA binding | 2.6 | 2.7E-02 |
| TE1 | MEGs | Lipid binding | 12.1 | 2.4E-02 |
| TE1 | MEGs + PEGs | Lipid binding | 15.5 | 2.8E-04 |
| TE1 | MEGs + PEGs | transcription factor activity, sequence-specific DNA binding | 2.5 | 1.5E-02 |
| All | MEGs | transcription factor activity, sequence-specific DNA binding | 2 | 3.7E-03 |
| All | MEGs | sequence-specific DNA binding | 3.3 | 5.5E-03 |
| All | MEGs | RNA polymerase II transcription factor activity, sequence-specific DNA binding | 3.3 | 3.6E-02 |
| All | MEGs | oxidoreductase activity, acting on the CH-OH group of donors, NAD or NADP as acceptor | 8.9 | 4.4E-02 |
| All | MEGs | lipid binding | 5.1 | 4.4E-02 |
| All | PEGs | poly(A) binding | 45.1 | 4.2E-02 |
| All | MEGs + PEGs | lipid binding | 5.7 | 3.9E-03 |
| All | MEGs + PEGs | sequence-specific DNA binding | 2.8 | 9.5E-03 |
| All | MEGs + PEGs | transcription factor activity, sequence-specific DNA binding | 1.8 | 9.7E-03 |
| All | MEGs + PEGs | RNA polymerase II transcription factor activity, sequence-specific DNA binding | 2.9 | 3.5E-02 |

**STable 3. Gene Ontology (GO) term enrichment analysis.** GO term enrichment analysis performed with DAVID Bioinformatics Resources version 2021 (Huang et al., 2009a; Huang et al., 2009b). Significantly enriched GO terms were determined using the EASE score (p-value < 0.05; (Hosack et al., 2003)).

| Study | MEGs | PEGs | Ecotype | Stage[b] | Temp | Extraction method | Contamination assesment |
|---|---|---|---|---|---|---|---|
| Pignatta et al. 2014 | 285 | 103 | Col, L*er*, Cvi | 6G | 21˚C | Manual dissection | Reciprocal crosses[f] |
| Hornslien et al. 2019 | 282 | 35 | Col, L*er*, Tsu | 4G | 18˚C | Whole seed | Reciprocal crosses[f] |
| Del Toro-De León et al. 2019 | 777 | 148 | Col, L*er* | 4LG/EH | 21˚C | INTACT[c] | Tissue enrichment analysis[g] |
| Picard et al. 2021[a] | 275 | 74 | Col, Cvi | 4G | 20˚C | FANS[d] | Reciprocal crosses[f] |
| This study | 181 | 56 | Col, Tsu | 4G/7LC | 20-22˚C | FANS[e] | Endosperm specific markers[h] |

**STable 4. Summary of experimental set-up of various imprinting analysis studies.** [a] Weak MEGs and PEGs identified by (Picard et al., 2021) were not included. [b] Number indicates at which day after pollination tissue was collected and letter indicates developmental embryo stage (G: globular, LG: late globular, EH: early heart, LC: late cotyledon). [c] INTACT method to purify endosperm nuclei (Moreno-Romero et al., 2017). [d] Fluorescence-activated nuclear sorting of 3C/6C endosperm nuclei stained with Partec CyStain UV Precise P nuclei staining buffer. [e] Fluorescence-activated nuclear sorting of endosperm nuclei with GFP-tagged histones. [f] In addition, seed compartment transcript profiles were analyzed (Belmonte et al., 2013). [g] Tissue enrichment test (Schon and Nodine, 2017). [h] Ploidy analysis of 3C/6C specific endosperm nuclei peaks.

| AGI | Forward primer | Reverse primer |
|---|---|---|
| AT1G02450 | AAAGGATCCCGGTCATCCAAAGAAACGAACAGTG | ATTCTCGAGTTAGAGAAAGTGATTGATTTTGGAGAGTGAAG |
| AT1G11190 | SAP COLLECTION | |
| AT1G17020 | TGTATAGAAAAGTTGCTCCAACTGGTCACTGTCGTGTTCAT | TTTTGTACAAACTTGGTCGGCTTCTTTCCTGGACGAAACA |
| AT1G25410 | SAP COLLECTION | |
| AT1G48930 | SAP COLLECTION | |
| AT1G62080 | SAP COLLECTION | |
| AT1G68510 | SAP COLLECTION | |
| AT2G25940 | SAP COLLECTION | |
| AT2G27380 | SAP COLLECTION | |
| AT2G34315 | SAP COLLECTION | |
| AT2G38900 | SAP COLLECTION | |
| AT2G41290 | AATGTCGACGCTGTATCCGGTTAGATATG | CGAATGGACCTTCATCCAATCCAC |
| AT2G42430 | ATAGAAAAGTTGGCGGAAGAACTTATAAAATAACTTTT | TGTACAAACTTGCGGCGAAACGAACAAAAAAGT |
| AT2G43660 | SAP COLLECTION | |
| AT3G12110 | AATGGATCCCAGTCGTACAAAGAAACTCTGCTTATAC | AATCTCGAGTCCAGCCTAAGGAATTGGAAAC |
| AT3G19660 | ATAGAAAAGTTGTATTTTCCAGAGCAGATATCGGAAT | TGTACAAACTTGCTTCTCCTTCTCCTTCTTGCTTCTC |
| AT3G22540 | TGTATAGAAAAGTTGTGTTGTTAGCAAGCATATCAAAAA | TGTACAAACTTGTTTCACTAAAGAAGCCTATAAGCT |
| AT3G29190 | SAP COLLECTION | |
| AT3G29770 | GTTGGATCCAAATATCAGTCGGGGTTACCG | AATCTCGAGTGCTTGAACATGAAGAAGAGTG |
| AT3G45130 | SAP COLLECTION | |
| AT3G57540 | SAP COLLECTION | |
| AT3G60245 | GGGGACAACTTTGTATAGAAAAGTTGTGGAACCATCTTTTGGGTTCCT | GGGGACTGCTTTTTTGTACAAACTTGCCACGCCGTCGTAGATGAGA |
| AT4G00180 | SAP COLLECTION | |
| AT4G00220 | ATAGGATCCCCCGCAAGTACAACCTCAGT | AAACTCGAGCTTCCTTTTCTACCACACAAGTCC |
| AT4G04460 | SAP COLLECTION | |
| AT4G05320 | AATGGATCCAGTCTAGCTCAACAGAGCTT | ATTCTCGAGCTGTTAATCAGAAAAACTCAGATTAATC |
| AT4G21980 | SAP COLLECTION | |
| AT4G30960 | AATGGATCCCCACAAGGACCACAACGAATG | AAGCTCGAGTCCGACCATGGTTTTTTACTTTAC |
| AT4G30970 | AGGGGATCCGTTTTCTCTGTTGCATTCCTCAATC | GGTCTCGAGTCGATGTTGCTAAGGCCAAAATGG |
| AT4G30970 long | GGGGACAACTTTGTATAGAAAAGTTGGAGTTTTCTCTGTTGCATTCCTCAATC | GGGGACTGCTTTTTTGTACAAACTTGGTGCATGTTGCTAAGGCCAAAATGG |
| AT4G31060 | SAP COLLECTION | |
| AT5G09370 | SAP COLLECTION | |
| AT5G10220 | GGGGACAACTTTGTATAGAAAAGTTGCTAATAACATTTGCAAGTTTTGTTTAAGA | GGGGACTGCTTTTTTGTACAAACTTGCTCTCCGACCACTGAATATTTCTCTA |
| AT5G13600 | GGGGACAACTTTGTATAGAAAAGTTGCTAGTTGAACTTCGAGGGTCCTAATAA | GGGGACTGCTTTTTTGTACAAACTTGCCTTTGTTCTTTCACTTCAACTCTTT |
| AT5G45890 | GGGGACAACTTTGTATAGAAAAGTTGTGTGTTCATGAGGTGATTGTGATT | GGGGACTGCTTTTTTGTACAAACTTGTGTTTTAGGAAAGTTAAATGACTTTTG |
| AT5G46950 | SAP COLLECTION | |
| AT5G50260 | SAP COLLECTION | |
| AT5G54650 | SAP COLLECTION | |
| AT5G56620 | SAP COLLECTION | |

**STable 5. Cloning primers of the endosperm-domain marker construct lines.** Primers to clone promoter sequences for candidate promoter genes were designed or constructs were obtained from the SAP collection (Benhamed et al., 2008).

| Domain | BR1 | BR2 | BR3 |
|---|---|---|---|
| EE | 1.341033 | 0.780990 | 1.026821 |
| ESR | 0.951849 | 0.977524 | 1.079423 |
| TE1 | 1.005221 | 1.463373 | 0.756521 |

**STable 6. Library normalization factor for marker line replicates.** Directly after mapping with bowtie2, library normalization factors for each biological replicate of EE, ESR and TE1 were determined by dividing the total read number of each replicate by the average number of reads across all replicates.
